# Single-cell profiling identifies a spectrum of human unconventional intraepithelial T lineage cells

**DOI:** 10.1101/2022.05.24.492634

**Authors:** Lore Billiet, Laurenz De Cock, Guillem Sanchez Sanchez, Rupert L. Mayer, Glenn Goetgeluk, Stijn De Munter, Melissa Pille, Joline Ingels, Hanne Jansen, Karin Weening, Eva Pascal, Killian Raes, Sarah Bonte, Tessa Kerre, Niels Vandamme, Ruth Seurinck, Jana Roels, Marieke Lavaert, Filip Van Nieuwerburgh, Georges Leclercq, Tom Taghon, Francis Impens, Björn Menten, David Vermijlen, Bart Vandekerckhhove

## Abstract

In the human thymus, a CD10^+^ PD-1^+^ TCRαβ^+^ differentiation pathway diverges from the conventional single positive T cell lineages at the early double positive stage. These cells are phenotypically and functionally similar to murine unconventional intraepithelial lymphocyte (uIEL) precursors. Here, the progeny of the human uIEL lineage was identified in antigen-inexperienced blood. The uIELs in thymus and peripheral blood share a transcriptomic profile, characterized by hallmark transcription factors (i.e. *ZNF683* and *IKZF2*), and polyclonal TCR repertoire with autoreactive features, exhibiting a bias towards early TCR alpha chain rearrangements. Single-cell RNA sequencing confirmed a common developmental trajectory between the thymic and peripheral uIELs, and clearly delineated this unconventional lineage in peripheral blood. This population is phenotypically defined as CD3^+^ TCRαβ^+^ CD4^-^ CCR7^-^ CD26^-^. It contains CD10^+^ recent thymic emigrants, Helios^+^ KIR^+^ CD8^+^ Tregs and CD8αα^+^ T cells. Thus, the uIEL lineage represents a well-defined but heterogeneous, unconventional TCRαβ^+^ lineage mostly confined in human within the CD8 single positive T cells.

**Summary:** Billiet et al. identify the postthymic progeny of the intraepithelial lymphocyte precursors in human based on shared characteristics of the T cell receptor repertoire and the transcriptome. This lineage represents a well-defined but heterogeneous, unconventional TCRαβ^+^ lineage mostly confined within the CD8 single positive T cells.

## INTRODUCTION

After successful rearrangement of the T cell receptor (TCR) alpha and beta locus in the thymus, each precursor T cell expresses a single, unique TCR. These cells are subsequently selected for major histocompatibility complex (MHC) binding affinity and only the cells with a moderate binding affinity further differentiate to mature CD4 or CD8 single positive (SP) conventional T cells (CTCs), a process called positive selection. Conventional T cells leave the thymus as dormant, stem cell-like cells without effector function. These cells will acquire effector function only after encountering their cognate foreign antigen. Concurrently, several minor lineages of so-called unconventional T cell populations are generated by different selection mechanisms. Immature thymocytes may receive a strong TCR signal, resulting in agonist selection. These T cells leave the thymus in an activated state, thereby facilitating rapid expansion in the tissues. Mucosal-associated invariant T (MAIT) and natural killer T (NKT) cells are two unconventional populations that express a semi-invariant TCRαβ reactive to microbial compounds. Besides these two oligoclonal UTC populations, an agonist-selected, polyclonal intraepithelial lymphocyte (IEL) lineage has been described ^1,2^.

CD8αα IELs form a prominent thymus-derived T cell population that guards the gut epithelium, in addition to conventional memory T cells ^3^. Their development has been predominantly studied in mice, where it was shown that thymic IEL precursors (IELps) diverge from the conventional T cell lineage upon high-affinity TCR interaction at the CD4^+^ CD8^+^ double positive (DP) stage. These precursor T cells are induced to differentiate into CD4^-^ CD8^-^ double negative (DN) T cells that express gut homing receptors ^4–6^. IELp TCRs were shown to exhibit broad reactivity against multiple MHC haplotypes and across MHC classes and therefore may be more generally deployable compared to CTCs that have a narrow antigen specificity ^7^. It has been reported in mice that IELps progress through a PD-1^+^ stage before upregulating T-bet, a process that is regulated by the transcription factors (TFs) *ID2, ID3* and *IKZF2* (Helios)^8^. These IELs may play a role in homeostasis by potentiating innate immune responses and/or by directly killing autologous infected cells. However, their mode of action currently remains enigmatic. It is currently not known whether CD8αα IELs present in other tissues or cancer micro-environments are all derived from the same agonist-selected T cell lineage ^9,10^.

CD8αα^+^ CD8β^-^ IELs are not present in the human gut, suggesting that there may not be a human equivalent for the murine CD8αα IEL lineage ^11^. However, it is possible that the human IELs have a different phenotype compared to their murine counterparts. Our research previously described the likely IELps in human postnatal thymus (PNT) that is similar to the murine thymic precursors of the CD8αα^+^ IEL lineage. These human IELps present in the thymus with a CD10^+^ PD-1^+^ phenotype and, unlike in mice, express both CD8αα and CD8αβ dimers ^12^. Similar to murine IELps, these cells can be generated *in vitro* by agonist stimulation of DP thymocytes, have a characteristic TCR repertoire different from CTCs, express TCRs with characteristics of autoreactivity and are activated in the thymus by high affinity ligands as evidenced by the expression of PD-1 and Helios ^12–14^. The existence of this lineage in the human thymus as well as the early divergence from the conventional T cell lineage was recently confirmed via single-cell mapping ^15,16^. It is currently unknown whether these human IELps leave the thymus and, if so, which phenotype and function these cells have.

The aim of this study is to identify the progeny of the IELp lineage in the human periphery. Within peripheral blood, the focus was on cord blood (CB) as CB CTCs have not yet been activated by foreign antigens and therefore, are easily discriminated from activated unconventional T cells. Therefore, the PNT IELp population and their likely progeny in CB were comprehensively analyzed by means of transcriptome, TCR and proteome analyses. A single-cell RNA sequencing (scRNA-seq) analysis was performed to further unravel the progeny in CB, combined with CITE-sequencing to establish a phenotypic definition encompassing the entire heterogeneous population. In summary, a detailed picture of the unconventional IEL (uIEL) lineage cells is provided in CB, which may lead to a better understanding of these cells in autoimmunity as well as immune reactivity to pathogens and will facilitate the study of the uIEL lineage in human.

## RESULTS

### The PNT and CB PD-1^+^ population share a similar transcriptomic and proteomic profile

The phenotype of the IELps in human PNT was defined as CD3^+^ TCRαβ^+^ CD4^-^ CD8α^+^ CD10^+^ PD-1^+^ (fig. S1A). The progeny of the PNT IELp population in human CB was tentatively defined as CD3^+/low^ TCRγδ^-^ CD4^-^ CD8α^+^ PD-1^+^ (fig. 1A)^12^. CD10 membrane expression, which is prominent on PNT PD-1^+^ IELps (fig. S1A), was less prominent on the CB PD-1^+^ population (fig. 1B). However, *MME* mRNA (encoding CD10) was significantly upregulated in the CB PD-1^+^ population compared to the conventional CD3^+^ PD-1^-^ population (fig. 1C). To examine the relatedness of PNT and CB PD-1^+^ populations, both were comprehensively analyzed by means of transcriptome and proteome analyses and compared to the CD3^+^ PD-1^-^ population. Principle component analysis (PCA) of the sorted CB populations indicated that the PD-1^+^ populations from the different donors clustered together and, similar to the respective PNT populations, shared more features with the unconventional TCRγδ population than with conventional PD-1^-^ populations (fig. 1D, S1C). Volcano plots comparing the transcriptomes of the PD1^+^ and PD1^-^ populations, showed significant upregulation in both CB and PNT PD-1^+^ populations of the hallmark TFs *ZNF683* (Hobit), *IKZF2* (Helios), *RUNX3, ID3* and *TBX21* (T-bet), and downregulation of *RORA* and *FOXP1* that mediates quiescence, and of *SATB1* that is required for positive and negative selection ^8,17,18^. Notably, both CB and PNT PD-1^+^ populations highly expressed the unconventional TCRαβ marker *TRGC2* (T Cell Receptor Gamma Constant 2), a gene progressively silenced in the conventional T cell lineage during passage through the CD4^+^ CD8^+^ DP stage in the thymus (fig. 1C, S1B)^19^. Mass spectrometry-based proteomics confirmed upregulation of Helios in the PNT PD-1^+^ population (fig. S1B) and, although not reaching the significance threshold, in CB (fig. 1C). Flow cytometric analysis validated the upregulation of Helios in both PNT and CB populations (fig. 1E, S1D). In search of additional distinctive markers, membrane proteins identified in the transcriptomic and proteomic profile of the PNT and CB PD-1^+^ populations, were confirmed by flow cytometry: CD3^low^ CD8β^low^ CCR7^-^ and EVI2B^+^ (fig. 1E, S1D). As expected, a correlation between the significantly differentially expressed genes and abundance of the corresponding proteins was observed (S1E). When zooming in on the genes or proteins that were differentially expressed between the PD-1^+^ and PD-1^-^ populations either in PNT or in CB, a highly significant positive correlation was revealed between the respective PNT and CB populations at both the RNA and protein level (fig. 1F, S1F, table S1). Supporting this correlation between the PNT and CB PD-1^+^ population, Gene Set Enrichment Analysis (GSEA) confirmed that the significantly upregulated genes in the PNT PD-1^+^ population were also significantly enriched in the CB PD-1^+^ population (fig. 1G). Based on the similarities in their transcriptomic and proteomic profile, it is hypothesized here that the CB CD3^+/low^ TCRγδ^-^ CD4^-^ CD8α^+^ PD-1^+^ population is the progeny of the PNT PD-1^+^ IELps.

**Fig. 1.**
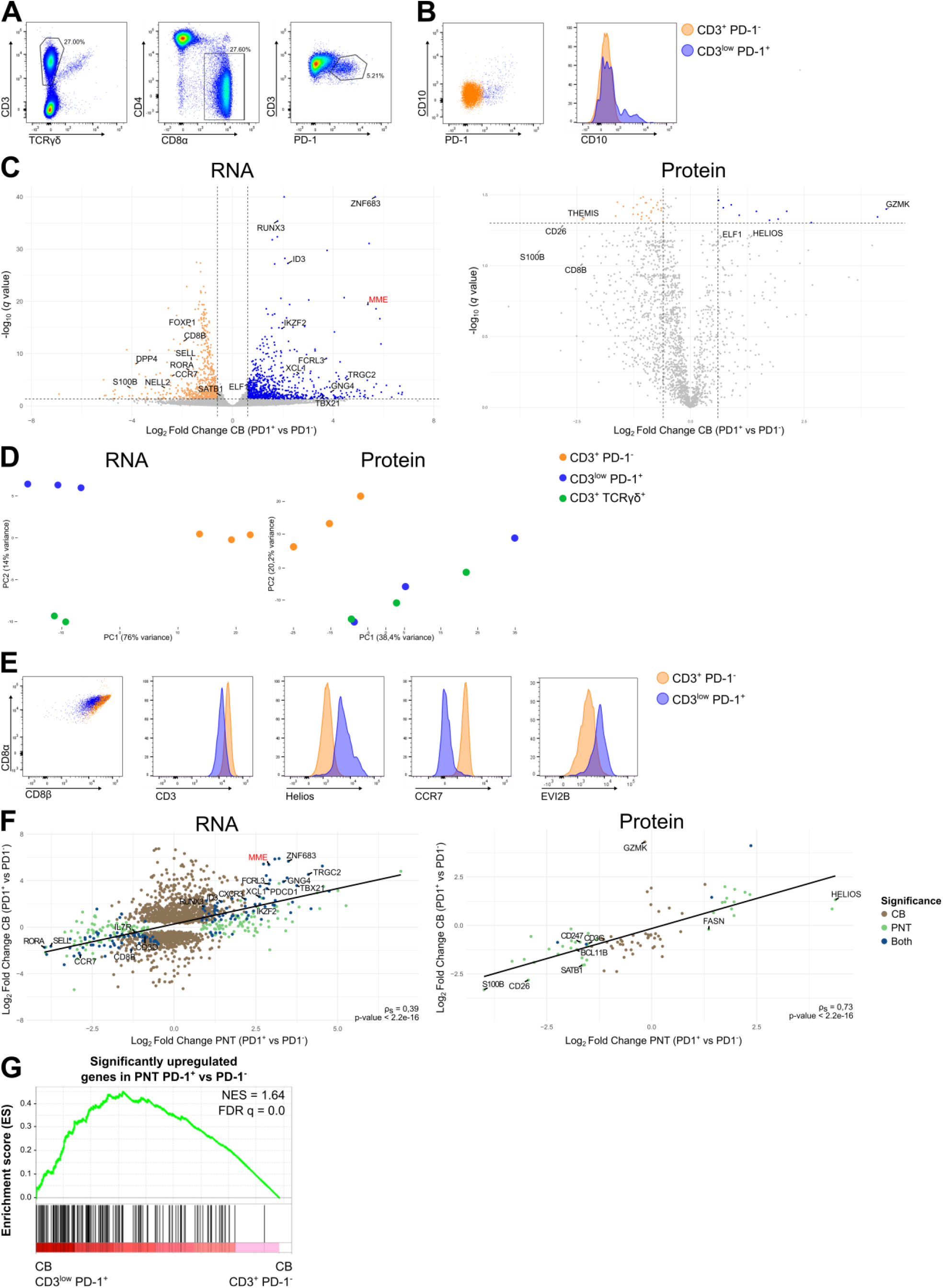
RNA and protein expression profile by the CB CD3^+/low^ PD-1^+^ population. (**A**) Representative gating strategy for the CD3^+/low^ TCRγδ^-^ CD4^-^ CD8α^+^ PD-1^+^ population in human CB. (**B**) CD10 expression on the CD3^+^ PD-1^-^ (orange) and CD3^+/low^ PD-1^+^ (blue) populations in CB, representative of at least three CBs. (**C**) Volcano plots of differentially expressed genes (left) and proteins (right) between the CB CD3^+/low^ PD-1^+^ and CD3^+^ PD-1^-^ populations. Triangles indicate data points outside the y-axis range. Data points with a Log_2_ Fold Change| > 0.6 and adjusted P < 0.05 are colored (upregulated in blue, downregulated in orange). (**D**) PCA of the transcriptome (left, donor corrected) and proteome (right) analysis of the sorted populations from three different CB donors. (**E**) Flow cytometric analysis of CD8α, CD8β, CD3, Helios, CCR7 and EVI2B on the CD3^+^ PD-1^-^ and CD3^+/low^ PD-1^+^ populations in CB, representative of at least three CBs. (**F**) Scatterplots and Spearman correlation coefficient comparing the Log_2_ Fold Change of the significantly differentially expressed genes (left) and proteins (right) of CB (CD3^+/low^ PD-1^+^ versus CD3^+^ PD-1^-^) and PNT (CD10^+^ PD-1^+^ versus CD10^−^ PD-1^-^). (**G**) GSEA showing the significantly upregulated gene set from PNT CD10^+^ PD-1^+^ versus CD10^−^ PD-1^-^, on the CB CD3^+/low^ PD-1^+^ versus CD3^+^ PD-1^-^ population. Normalized enrichment score (NES) and false discovery rate q value (FDR q) are shown.

### The TCR repertoires of the PNT PD-1^+^ IELp and CB PD-1^+^ populations share highly characteristic features

TCRα rearrangements are known to occur in a non-random manner, starting at the J-proximal V segments and the V-proximal J segments. As multiple sequential rearrangements may occur during the DP thymocyte stage, later rearrangements tend to be biased towards J-distal V segment and V-distal J segment usage. As shown in our previous publication and confirmed by others, the TCRα usage of PNT PD-1^+^ IELp population is biased towards early rearrangements similar to early DP thymocytes, in contrast to late DP and conventional thymocytes ^12,13,15^. Additionally, an association between autoreactive populations (T_reg_ and IELp) and CDR3 amino acidic (AA) residue content, including the presence of cysteines within 2 positions of the CDR3 apex (cysteine index) and enrichment of hydrophobic AA doublets at positions 6 and 7 of the CDR3 (hydrophobic index), is reported ^13,20^. Similar CDR3 properties are also reported to result in strong TCR-ligand interactions ^21–23^. Finally, in contrast to NKT or MAIT cells, the repertoire of PNT IELps was determined to be polyclonal (fig. S2A-B). Thus, to obtain additional evidence for a precursor-progeny relationship between the PNT and CB PD-1^+^ populations, the TCR repertoire of the CB PD-1^+^ population was analyzed for these characteristics and compared to the PNT PD-1^+^ TCR repertoire (fig. S2A-F for PNT).

Analysis of the CDR3α and CDR3β clonotypes revealed that the CB PD-1^+^ population was polyclonal and the degree of polyclonality was similar to the conventional PD-1^-^ T cell population (fig. 2A). This was quantified by calculating the D75 values, i.e. the percentage of unique clonotypes required to occupy 75% of the total TCR repertoire, for both populations. The D75 values were about 30% for both CB populations, indicating that the bulk of the repertoire consists of a wide variety of clonotypes. Moreover, no significant mean difference between the two CB populations could be identified, supporting the notion that the unconventional T cell population was equally polyclonal as the conventional T cell population in CB (fig. 2B). The CB PD-1^+^ population exhibited biased usage of early J-proximal TCR Vα rearrangements and early V-proximal TCR Jα rearrangements, and this bias was similar to that found in the PNT PD-1^+^ IELps (fig. 2C-E, S2D-E). The cysteine index of the TCRβ chain was significantly higher in the CB PD-1^+^ population compared to the PD-1^-^ population and the same trend could be observed for the TCRα chain (fig. 2F). The hydrophobic index of the TCRβ chain was likewise significantly higher in the CB PD-1^+^ population compared to the PD-1^-^ population. However, such a trend could not be established for the TCRα chain (fig. 2G). In line with this, the CDR3β repertoire exhibited higher interaction strength values (strength and volume parameters) compared to the PD-1^-^ population counterpart (fig. 2H)^21,22^. In contrast, polarity, a property associated with conventional T cells, was reduced in the CB PD-1^+^ population (fig. 2H)^24^. Finally, TCR sequencing of the CB populations revealed a significant higher percentage of TRAJ sequences using the T Cell Receptor Delta Variable 1 (*TRDV1*) gene segment (instead of a TRAV gene segment) in the CB PD-1^+^ population (fig. 2I). When subsequently analyzing the PNT populations, an increased *TRDV1* usage was also observed in the PNT PD-1^+^ IELps compared to their PD-1^-^ counterparts (fig. S2H). Vδ1^+^ cells expressing a hybrid TRDV1-TRAJ-TRAC TCR chain and co-expressing a TCRβ chain rather than a TCRγ chain have been previously reported in human peripheral blood. This population, termed δ/αβ T cells, recognizes antigens presented by both human leukocyte antigen (HLA) and CD1d ^25^. By using an anti-Vδ1 antibody, an enrichment of Vδ1 membrane expression was shown in the CB PD-1^+^ population (fig. 2J, S2I).

**Fig. 2.**
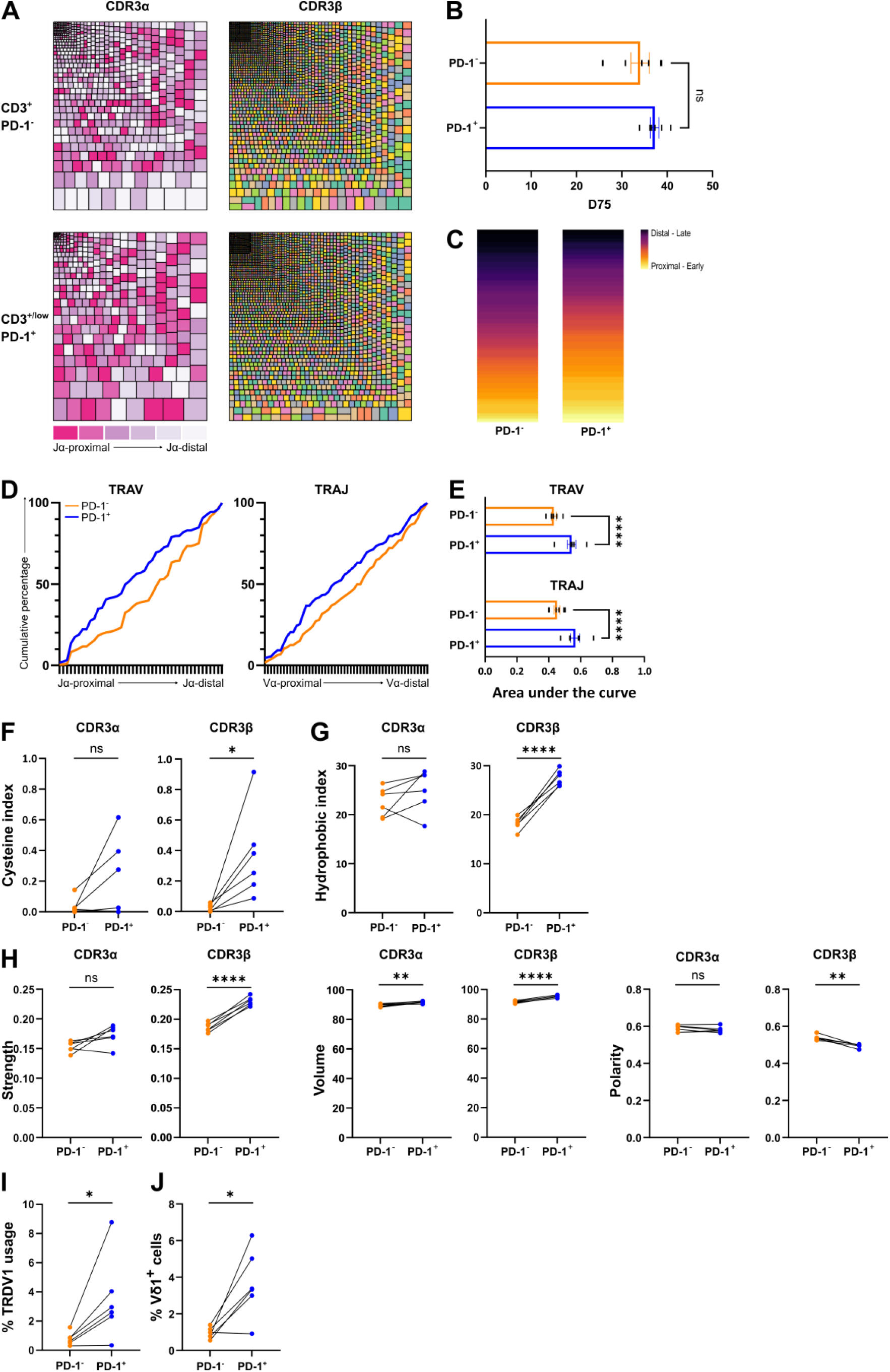
Distinctive TCR repertoire of the CB PD-1^+^ population. (**A**) Representative tree maps showing CDR3α (left) and CDR3β (right) clonotype usage in relation to repertoire size for the CD3^+^ PD-1^-^ (top) and CD3^+/low^ PD-1^+^ (bottom) populations. Each rectangle represents one CDR3 clonotype and its size corresponds to its relative frequency in the repertoire. Rectangle colors for CDR3α are categorized from J-proximal (pink) to J-distal (white) for the TRAV gene segments and for CDR3β are chosen randomly. (**B**) D75 (percentage of clonotypes required to occupy 75% of the total TCR repertoire) analysis comparing the PD-1^-^ and PD-1^+^ population (individual values and mean ± SEM). Mean difference was not significant. (**C**) Representative heatmap illustrating the difference in J-proximal versus J-distal TCR Vα usage between the PD-1^-^ and PD-1^+^ populations. (**D**) Cumulative percentage of TRAV (left) or TRAJ (right) gene segment usage by the PD-1^-^ (orange) and PD-1^+^ (blue) population in a representative donor. X-axis represents the location in the TRAV or TRAJ locus. (**E**) Area under the curve determined from the cumulative plots from each sample (individual values and mean ± SEM). Šídák’s multiple comparisons test was used to assess the statistically significant difference. p-value < 0.0001 (****). (**F**) Cystein index (percentage of unique sequences with cysteine within 2 positions of the CDR3 apex) and (**G**) Hydrophobic index (percentage of unique sequences with self-reactive hydrophobic CDR3 position 6 and 7 doublets) of the CDR3α (left) and CDR3β (right). (**H**). Physicochemical properties (strength, volume and polarity) of the CDR3α (left) and CDR3β (right). (**I**) Percentage of unique sequences containing a *TRDV1* segment. (**J**) Flow cytometric analysis of the percentage of Vδ1^+^ (A13 clone) cells in both CB populations. (**B, F-I**) Paired t-tests were used to assess statistical significance. Connected values correspond to paired populations of the same biological replicate (n= 6). p-value > 0.05 (ns), p-value < 0.05 (*), p-value < 0.0001 (****).

To conclude, a series of characteristic features of the TCR repertoire of the PNT PD-1^+^ IELp can be tracked within the CB PD-1^+^ population. This strongly suggests that the CB PD-1^+^ T cell population is the progeny of the PNT PD-1^+^ T cell population and that biased TCRα chain usage and self-reactive features of both TCR chains are acquired during early thymic agonist selection and preserved after thymic egress.

### The CB unconventional T cell population extends beyond the CD3^+/low^ PD-1^+^ cells

Because murine IELps were shown to downregulate PD-1 expression during differentiation and PD-1^-^ (T-bet^+^) IELp populations were described ^26,27^, the unconventional CD3^+/low^ PD-1^+^ CB population may not include the complete human uIEL population, but only the more recent thymic emigrants of that population. Therefore, the discriminatory markers identified above were used in a flow cytometric analysis of the entire CD3^+/low^ TCRγδ^-^ CD4^-^ CD8α^+^ fraction in CB (fig. 3A). In addition to the expected conventional CCR7^+^ population, the resulting uniform manifold approximation and projection (UMAP) revealed a distinct cluster of CCR7^-^ cells that contained all PD-1^+^ cells (fig. 3A). Within this CCR7^-^ cluster, the majority of the PD-1^-^ cells were Helios^+^ and EVI2B^+^, two markers associated with the IELp population (fig. S1C). Here, it was hypothesized that these Helios^+^ and EVI2B^+^ cells may contain PD-1^-^ uIELs. Consequently, the CB uIEL population was further studied within the CCR7^-^ EVI2B^+^ as well as within the CD3^+/low^ PD-1^+^ population.

**Fig. 3.**
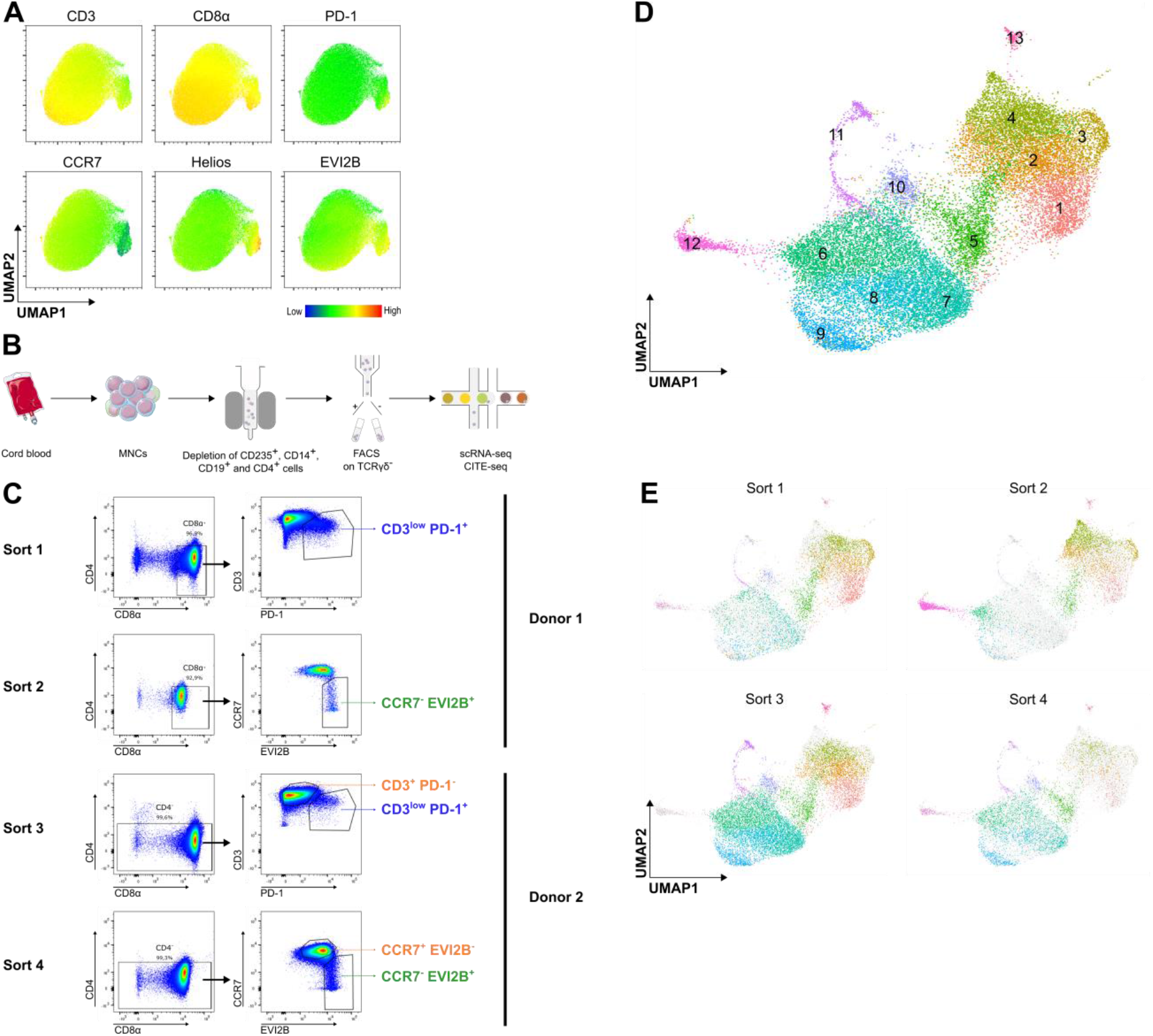
Single-cell RNA sequencing of human CB reveals a heterogeneous unconventional population. (**A**) UAMP analysis of flow cytometry data gated on all CD3^+/low^ TCRγδ^-^ CD4^-^ CD8α^+^ cells from CB. (**B**) Schematic workflow of the CB processing. (**C**) Gating strategy on the CD3^+/low^ TCRγδ^-^ cells for the four different sorts, to isolate the different populations of interest. (**D**) UMAP of 24 727 CB single cells, colored by the 13 identified cell clusters. **(E)** UMAP representation of the four different sorts, with the cells from the particular sorts colored according to the cell clusters and the remaining cells in grey.

To comprehensively study the heterogeneity of the uIELs in CB, single-cell RNA sequencing (scRNA-seq) was performed. CB of two different donors was depleted of the CD4^+^, CD14^+^, CD19^+^ and CD235^+^ cells (fig. 3B). Of the first donor, the uIELs were sorted as CD8α^+^ CD3^+/low^ PD-1^+^ (sort 1) or CD8α^+^ CCR7^-^ EVI2B^+^ (sort 2) within the CD3^+/low^ TCRγδ^-^ CD4^-^ window. Both fractions were labeled with different hashtags before they were further processed, enabling subsequent assignment of the single cells to their corresponding sorting strategy. For the second donor, the sorting strategy was expanded to include CD4^-^ CD8α^-^ DN cells (sort 3 and 4). In sort 3 and 4, the respective conventional populations were also sorted and added in equal portions before further analysis (fig. 3C). Sort 4 was combined with cellular indexing of transcriptomes and epitopes (CITE)-seq to capture expression of 277 membrane proteins. Using a droplet-based single-cell platform 3′ gene expression libraries were constructed. Reciprocal PCA (RPCA) was used to integrate the Seurat objects resulting from the separate sorts. This approach resulted in 24 727 cells included in the scRNA-seq analysis after quality control and filtering. Leiden clustering applied to this filtered and integrated Seurat object defined 13 distinct clusters (fig. 3D). Both CD8α^+^ CD3^+/low^ PD-1^+^ (sort 1) and CD8α^+^ CCR7^-^ EVI2B^+^ (sort 2) consisted mainly of clusters 1-5, suggesting that these clusters represent the unconventional T cells. Focusing on these 5 clusters, cluster 1 is relatively overrepresented in the CD3^+/low^ PD-1^+^ sorts 1 and 3 and cluster 4 is relatively enriched in the CCR7^-^ EVI2B^+^ sorts 2 and 4, highlighting that indeed the two different sorting strategies captured slightly different populations (Fig 3E).

### Defining unconventional T cell clusters using transcriptomics

This heterogeneity consisting of 13 different clusters has not been reported before for CB CD8^+^ T cells. Therefore, the clusters were manually annotated based on prominently upregulated genes (fig. 4A-B; table S2-3). One non-T cell cluster was detected, which was annotated as NK cells based on high expression of NK-associated genes (*GZMB, TYROBP*) and absence of membrane CD3 and TCRαβ (fig. 4A, S4A). The NK cluster was assumed to be a contaminant due to lenient gating for CD3. Two minor T cell clusters were annotated: an NKT/MAIT cluster based on high *KLRB1* expression and a cycling T cell cluster based on upregulated effector (i.e. *GZMA, GZMK*) and cycling genes (i.e. *PCNA, MKI67, CDC6*) (fig. 4A, S3A). The NKT/MAIT cells were the main “contaminant” in the CCR7^-^ EVI2B^+^ sorts (fig. S3B). Based on their upregulation of hallmark CD8αα^+^ and IELp T cell genes ^8,15^, five unconventional T cell (UTC) clusters were annotated (fig. 4A-B). The unconventional marker genes *TRGC2, PDCD1* and *IKZF2* were expressed homogeneously in these UTC clusters, with some expression in the cycling T and MAIT/NKT cells (fig. 4C). The remaining clusters were considered conventional T cell (CTC) clusters based on the expression of typical conventional naive T cell markers (*FOXP1, SELL*)(fig. 4A-B). Furthermore, markers (i.e. *NELL2, S100B*) which were strongly overexpressed in the bulk transcriptome of the conventional PD-1^-^ populations in both PNT and CB (fig. 1C, S1C), were homogeneously expressed in the CTC clusters and largely absent in the UTC clusters, except for the GZMK^+^ DN UTC cluster. Likewise, *DPP4* (encoding CD26) was expressed homogeneously and exclusively by the CTC clusters and strongly in the NKT/MAIT cluster (fig. 4C).

**Fig. 4.**
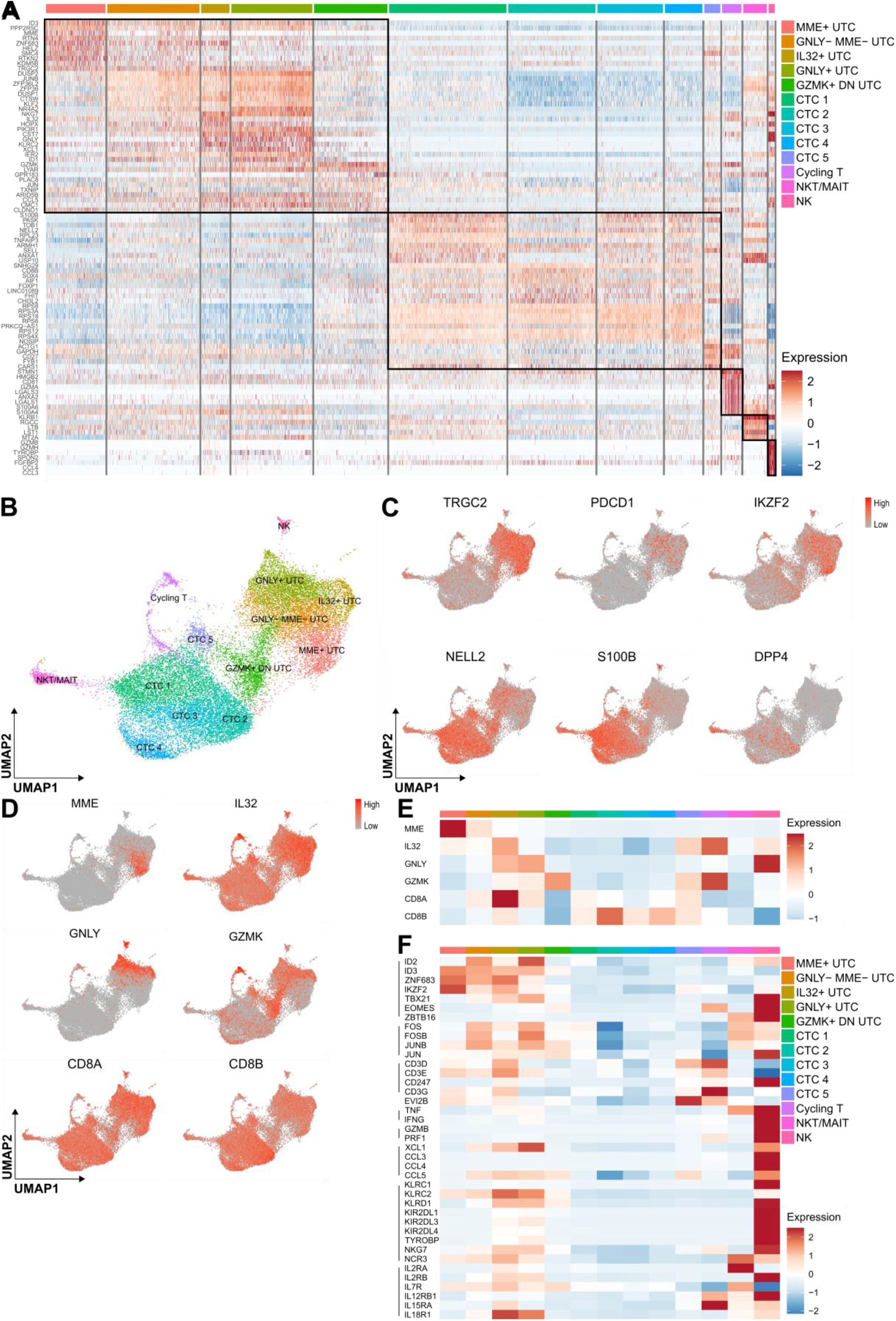
Annotation of the UTC clusters in CB. (**A**) Heatmap showing the expression of the top 10 differentially expressed genes per defined cluster in CB. Recurrent genes are not repeated. The genes are listed in Table S3. (**B**) UMAP visualization of the 13 identified cell clusters in CB. (**C**) UMAP feature plots representing discriminating genes between the UTC and CTC clusters. (**D**) UMAP feature plots representing differentially expressed genes used to annotate the different UTC clusters. (**E**) Heatmap showing the mean expression of the UTC signature genes in the different clusters in CB. (**F**) Heatmap showing the mean expression of characteristic UTC genes in the different clusters in CB.

Based on the differentially expressed genes, unique distinctive annotations were provided for the individual UTC clusters (table S3). *MME* and *GNLY* (encoding granulysin) are solely expressed by the UTC clusters. *IL32* is highly expressed by IL32^+^ UTCs but is also expressed at lower levels by different CTC and UTC clusters. The cluster that bridges the bulk of the UTCs and the CTCs had a characteristically high expression of *GZMK* without overexpression of other cytolytic effector genes and a low expression of *CD8A* and *CD8B*, and was annotated as the GZMK^+^ DN UTC cluster (fig. 4D-E).

The UTC clusters expressed the typical TFs of the IELp lineage, *ID3, ZNF683, IKZF2, RUNX3* and *TBX21*, although very limited in the GZMK^+^ DN UTCs. Importantly, expression of these TFs was absent in the other T cell clusters (fig. 4F). *ID3, ZNF683* and *IKZF2* were expressed highest in the MME^+^ UTCs. As expected, the CD3^+/low^ PD-1^+^ population (sort 1 and 3) was enriched for these MME^+^ UTCs, the recent thymic emigrants (fig. S3B). The activator protein 1 (AP-1) TFs (i.e. *FOS, JUN*) were constitutively expressed in both the UTC and NKT/MAIT clusters (fig. 4E). With regard to effector function, low constitutive expression of cytokines (i.e. *TNF* and *IFNG*) and genes involved in cytolysis (i.e. *GNLY*, granzymes) were observed in the GNLY^+^ UTCs, IL32^+^ UTCs and GZMK^+^ DN UTCs (fig. 4D-E). These three clusters were considered effector UTC clusters. These effector clusters showed expression of multiple NK receptors (i.e. *KLRC2, KLRD1, NCR3, KIRs*). *EOMES*, which is reported to be absent in murine uIELs, was weakly expressed in the effector clusters. Finally, UTCs expressed important components of the IL-2, IL-7 and IL-15 receptors, as well as IL-12 and IL-18 receptor components, which are required for inflammation-induced cytokine responses. Expression of the genes discussed above was mostly absent in CTCs and cycling T cells (fig. 4E).

### CB UTCs originate from ZNF683^+^ CD8αα^+^ thymocytes

It was established above that the TCR repertoires of the PNT PD-1^+^ IELp and CB PD-1^+^ populations are similar, suggesting that the thymic population is the precursor of the CB uIELs. To further explore this developmental pathway, the CB scRNA-seq dataset was integrated with a previously published PNT CD3^+^ scRNA-seq dataset ^15^, followed by a trajectory analysis. As published, the CD3^+^ DP PNT cells diverged into two main pathways: the unconventional pathway consisting of GNG4^+^ CD8αα(I), ZNF683^+^ CD8αα(II) and TCRγδ cells and the conventional pathway of CD4^+^ and CD8^+^ SP T cells (fig. 5A). Integration with the CB populations showed that the CB UTC clusters partially overlapped with the PNT UTCs, whereas the CB CTC partially overlapped with the PNT SP T cells (fig. 5A-B). When focusing on the UTC pathway, the CB MME^+^ UTCs overlapped with the PNT CD8αα(I) as well as CD8αα(II) (fig. 5B). Based on the expression of hallmark differentially expressed genes (i.e. *GNG4, MME* and *ZNF683*), the CB MME^+^ UTCs seemed to originate from the PNT ZNF683^+^ CD8αα(II) rather than from the GNG4^+^ CD8αα(I) (fig. 5C-D). Therefore, a TSCAN trajectory analysis was performed with the ZNF683^+^ CD8αα(II) as the population of origin. This analysis revealed a common pathway passing through PNT ZNF683^+^ CD8αα(II) and CB MME^+^ UTCs, leading to a branching point at the GNLY^-^ MME^-^ UTC cluster. The GNLY^-^ MME^-^ UTC cluster gave rise to three distinct lineages: the GNLY^+^ UTCs (lineage 1), the GZMK^+^ DN UTC (lineage 2) and the IL32^+^ UTCs (lineage 3)(fig. 8E-F). The common pathway included ZNF683^+^ cells, which upregulated *ID3* and differentiated in *TBX21*(T-bet) positive cells (fig. 5G-H). During terminal differentiation, all three lineages expressed high levels of AP-1 TFs and gradually upregulated different effector markers. When analyzing the data in regulons, the transcriptional regulation of the UTCs and the CTCs was significantly different. As expected, the CTCs were mainly regulated by the conventional TFs *FOXP1* and *RORA*, while in the UTCs, *KLF4*, which negatively regulates TCR-mediated proliferation in CD8^+^ T cells, and *RUNX3* were prominent (fig. 5I)^28,29^.

**Fig. 5.**
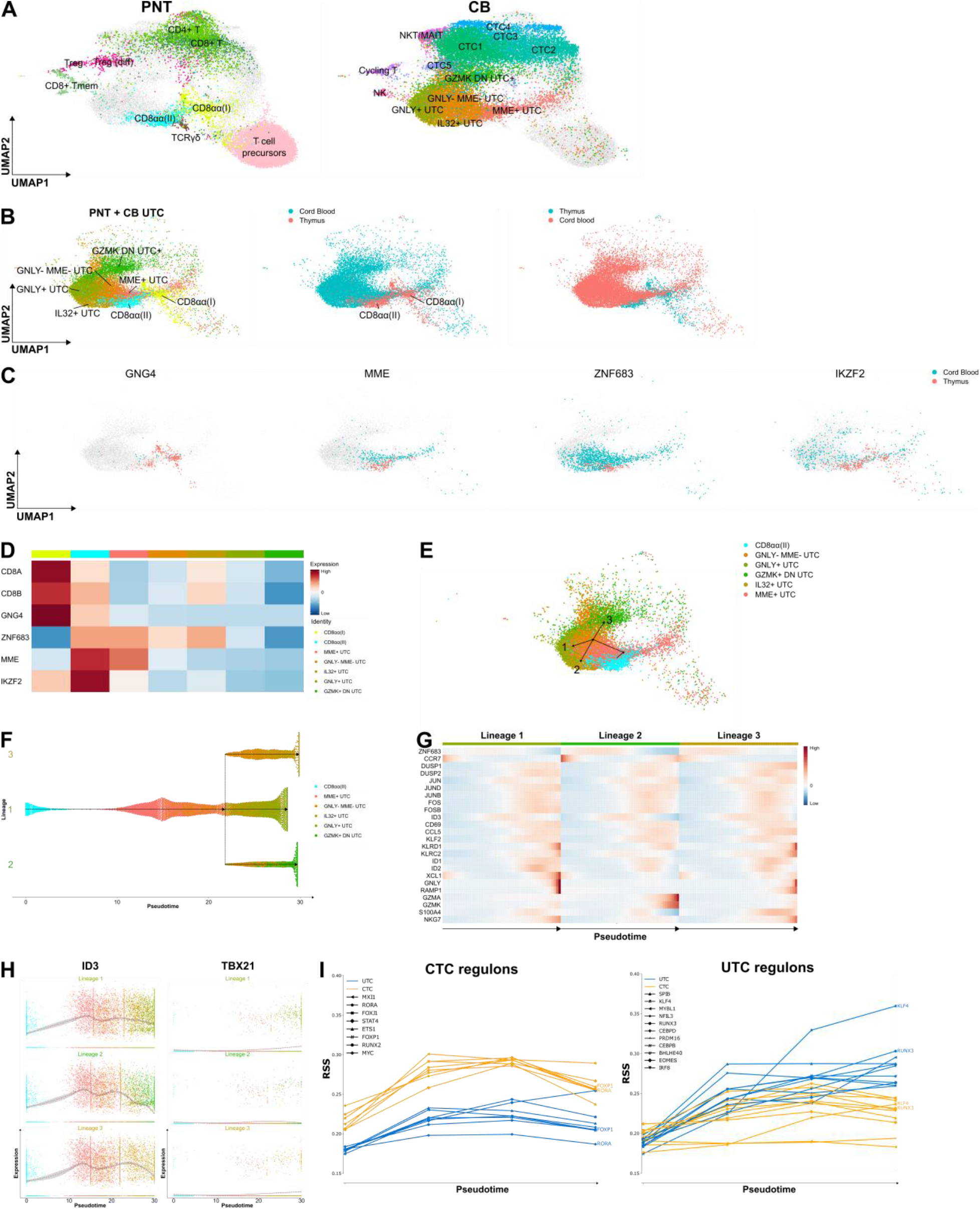
UTC pathway analysis reveals 3 effector lineages. (**A**) UMAP visualization of the annotated PNT clusters (left) and CB clusters (right) after integration. (**B**) UMAP visualization highlighting the PNT and CB UTCs and showing the overlay of the integrated UTCs derived from PNT or CB. (**C**) UMAP visualization *GNG4, MME, ZNF683* and *IKZF2* expression by UTCs from PNT (red) or CB (blue). Only cells with a relatively high expression are colored. (**D**) Heatmap showing the mean expression for hallmark differentially expressed genes per UTC population. (**E**) UMAP visualization of the TSCAN trajectory analysis of the UTC populations with the PNT ZNF683^+^ CD8αα(II) as the origin population. (**F**) Dendrogram of the predicted UTC lineages. (**G**) Heatmap showing the varying gene expression in the pseudotime for the different lineages. (**H**) For each lineage, *ID3* or *TBX21* expression is shown per single cell and summarized as the mean (grey line). **(I)** The regulon specificity score (RSS) is plotted per cell type for the most prominent UTC and CTC pathway. The cell populations on the x-axis are ordered according to pseudo-time.

### TCRαβ^+^ CCR7^-^CD26^-^cells represent the uIEL population in cord blood

To identify phenotypical differences between the different clusters, CITE-seq was included in sort 4 of our scRNA-seq experimental setup (fig. 3C). Membrane protein data were acquired for 3 615 single cells across all clusters (fig. 6A). CD10 (encoded by *MME*) was expressed exclusively in the MME^+^ UTC cluster. A small fraction of the MME^+^ UTC cluster expressed the thymic T cell immaturity marker CD1a, confirming that this cluster included the recent thymic emigrants. Likewise, PD-1 was expressed by the immature UTCs, as well as by the cycling T cells. In contrast, CD26 (*DPP4*) was strongly expressed by the CTCs and NKT/MAIT cells. The latter also highly expressed CD161 (*KLRB1*), similar to the NK cells. CD54 (*ICAM-1*) and CD244 (*2B4*) are known to be induced in many immune cell types during inflammatory responses ^30,31^. CD54 and CD244 were highly expressed by the effector type UTCs, but not by the earliest CD1a^+^ MME^+^ cells and the GZMK^+^ DN UTC cluster. Expression of NK receptors such as CD158b (*KIR2DL/DL3*), CD244 and CD94 was mainly observed in the GNLY^+^ UTC (fig. 6B, S4A, table S4).

**Fig. 6.**
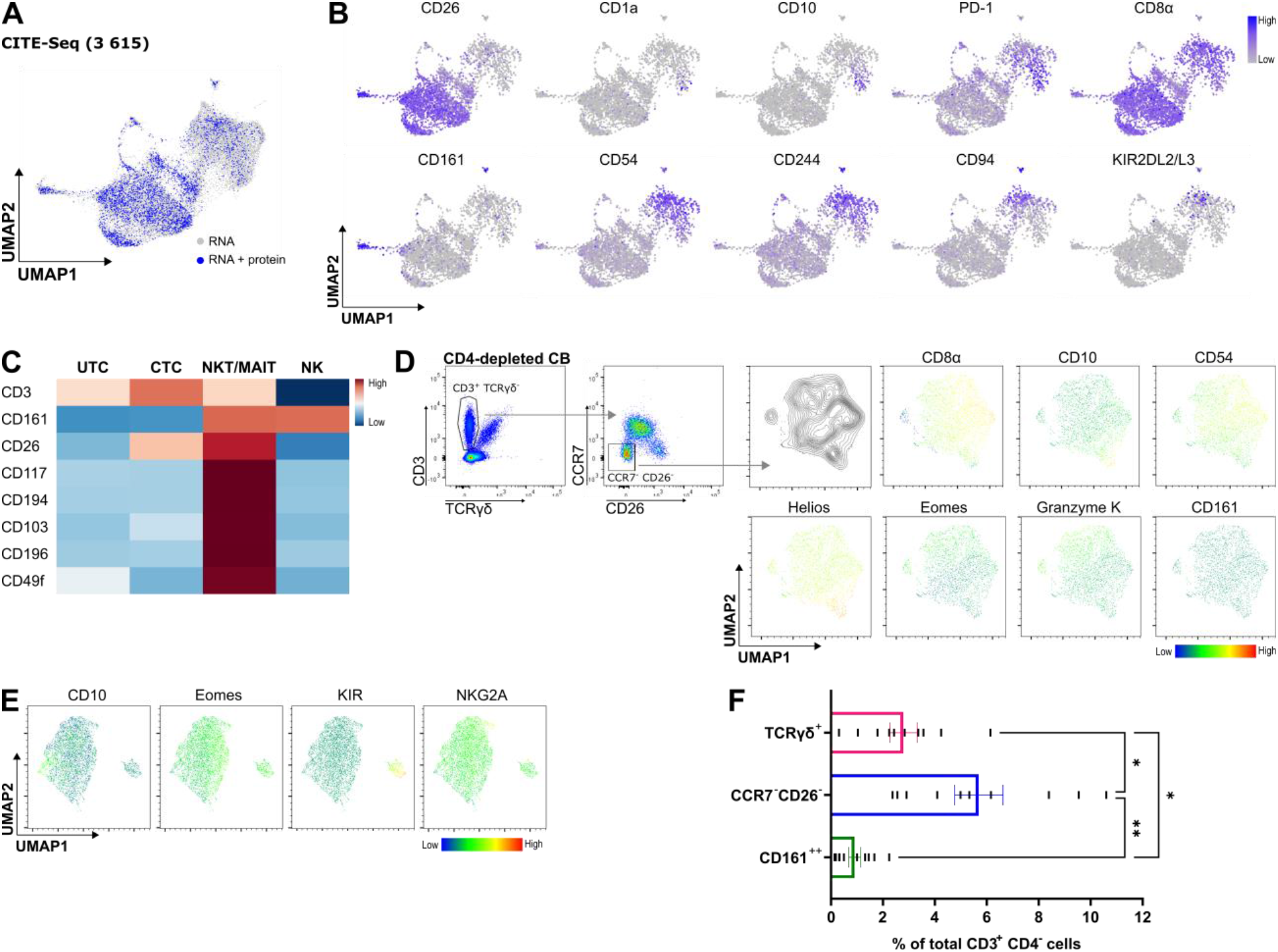
CCR7^-^CD26^-^constitutes the largest unconventional population in CB. (**A**) UMAP visualization of the single cells in the scRNA-seq analysis from which RNA data (grey) or combined RNA and protein data (blue) was determined. (**B**) Protein-based UMAP visualizations showing the expression of the indicated cell surface protein markers, only visualizing the single cells from which protein data was collected. (**C**) Heatmap depicting the mean expression of cell surface protein markers used to differentiate the UTC and CTC clusters from the NKT/MAIT and NK clusters. (**D**) Representative flow cytometric analysis of a CD4-depleted CB. The CD3^+/low^ TCRγδ^-^ cells were further gated for CCR7^-^ CD26^-^ cells, for which UMAP visualizations are shown. (**E**) Representative flow cytometric analysis of CD10, Eomes, KIR (KIR2DL1/DS1, KIR2DL2/DL3 and KIR3DL/DS1) and NKG2A expression on the CD3^+/low^ TCRγδ^-^ CD4^-^ CCR7^-^ CD26^-^population in CB, for which UMAP visualizations are shown. (**F**) Percentage of the CD3^+^ CD4^-^ cells in CB which are TCRγδ^+^, CCR7^-^ CD26^-^ or CD161^high^ (individual values and mean ± SEM, n = 10). Holm-Šídák’s multiple comparisons test was used to assess the statistically significant difference. p-value < 0.05 (*), p-value < 0.01 (**).

Because of their scarcity in human CB, the unconventional semi-invariant NKT/MAIT populations have only been studied to a limited extent. The NKT/MAIT cells are the main non-UTC “contaminant” in CCR7^-^ EVI2B^+^ cells (sort 2, fig. 3F). However, they can easily be discriminated from the UTCs using the markers CD161, CD26, CD117 (*KIT*), CD194 (*CCR4*), CD103 (*ITGAE*) and CD196 (*CCR6*) (fig. 6C). Our scRNA-seq and CITE-seq data did not incorporate TCR sequencing. Therefore no further distinction was made between NKT or MAIT cells.

Based on these CITE-seq results, the uIEL lineage in CB was redefined to include the effector UTC clusters. Flow cytometric analysis of the CD3^+^ TCRγδ^-^ cells of a CD4-depleted CB showed a distinct CCR7^-^ CD26^-^ population. The UMAP visualization of this CCR7^-^ CD26^-^ population clearly showed a gradient of Helios expression with the CD10^+^ cells having the highest expression. Whereas CD10 stained the immature UTCs, CD54 preferentially stains the effector UTCs. Finally, Eomes^+^ Granzyme K^+^ cells are included of which a minority is ultimately CD8α^-^. CD161^+^ NKT/MAIT cells are not included in the CCR7^-^CD26^-^ population (fig. 6D).

A population of virtual memory T (T_VM_) cells has been documented in mice, expressing TCRs with a strong binding affinity towards self-antigens. These cells express Eomes in the thymus and acquire a memory phenotype in the periphery ^32–34^. A population of CD8^+^ KIR/NKG2A^+^ T cells expressing *EOMES* has been described in CB and is proposed as a putative human analog of the T_VM_ population in mice ^35^. As KIR expression in CB T cells could only be observed in the UTC clusters (fig. 6B), the CD3^+^ TCRγδ^-^ CD4^-^ CCR7^-^ CD26^-^ population was further assessed for these markers. Flow cytometric analysis indeed showed that the cells with the highest expression for KIR or NKG2A expressed Eomes. scRNA-seq analysis clearly indicated that *EOMES, KIR, KLRC1* (encoding NKG2A) and *IL2RB* (encoding CD122) were, besides in the NK cluster, almost exclusively present in the UTC clusters and mainly enriched in the GNLY^-^ UTC cluster (fig. S4B). No separate cluster of T cells expressing these markers could be observed, suggesting that these T_VM_ cells are part of the uIEL population in human CB.

Finally, the size of the CD3^+^ TCRγδ^-^ CD4^-^ CCR7^-^ CD26^-^ population expressed as percentage of the CD3^+^ CD4^-^ T cells was determined (fig. 6F, S4C). Although the percentages in the different donors varied substantially, the uIEL population is generally significantly larger than the TCRγδ^+^ or CD161^high^ NKT/MAIT population in CB. It constitutes the largest unconventional T cell population in CB.

### Downregulation of CD8β is a unique characteristic of the CCR7^-^CD26^-^population

Many characteristic markers of the uIELs (i.e. PD-1, Helios) are related to activation by autoantigens in the thymus. Therefore, the stability and specificity of these hallmarks was tested during culture-expansion. Both CB populations were culture-expanded with interleukins only, as an *in vitro* equivalent for steady state persistence of the cells in tissues. To include the entire population, the CTCs were isolated from CD4-depleted CBs as CD3^+^ TCRγδ^-^ CD8α^+^ CCR7^+^ and the UTCs as CD3^+^ TCRγδ^-^ CCR7^-^ CD26^-^. Similar to the PNT PD-1^+^ population, the CB UTCs extensively proliferated in the presence of interleukin-15 (IL-15)(fig. S5A-B)^36^. CD26 expression was upregulated by the UTCs in culture, while Helios proved to be a stable distinguishing feature between the CTCs and UTCs, even after proliferation (fig. 7A, fig. S5C). CD158b (KIR2DL2/DL3) was expressed on a minority of the IELs and this expression remained stable during culture. Moreover, no expression could be observed on IL-7 culture-expanded CTCs (fig. 7A, S5D-E). This suggested that all KIR^+^ T cells detected *in vivo* belong to the UTC lineage. CD8β expression decreased on UTCs during culture. whereas expression on CTCs was stable (fig. 7B).

**Fig. 7.**
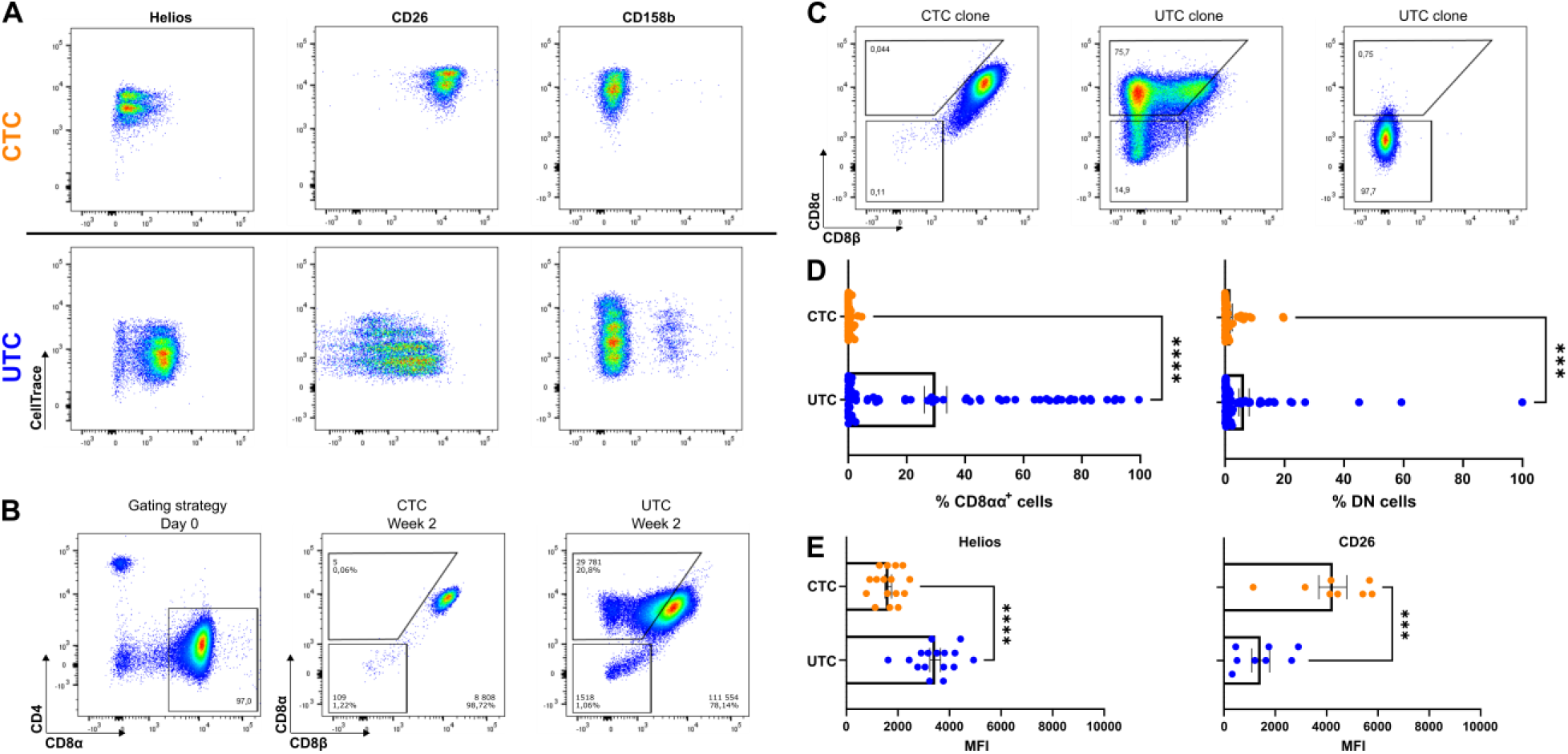
Stable phenotypic uIEL markers following proliferation. (**A**) Representative dot plots of flow cytometric markers expressed by the CTCs after 5 days of incubation with IL-7 (10 ng/mL, upper row) and by the UTCs after incubation with IL-15 (10 ng/mL, bottom row). Proliferation assessed by CellTrace Violet dye dilution. (**B**) Both the CTCs and UTCs were sorted as strictly CD4^-^ CD8α^+^ (left). Representative assessment of CD8α and CD8β expression by the CTC and UTC populations after two weeks of proliferation with IL-15. (**C**) Representative examples of CD8α and CD8β expression by CTC and UTC clones. (**D**) Percentage of CD8αα^+^ or DN cells (gated as shown in figure 7C) in CTC or UTC clones after at least 2 weeks of expansion. Individual values and mean ± SEM of 60 clones per population are shown. Mann-Whitney was used to assess statistical significance. (**E**) Mean fluorescence intensity (MFI) of Helios and CD26 expression by CTC or UTC clones. Individual values and mean ± SEM of 8 or 16 clones per population are shown. Unpaired t-tests were used to assess statistical significance. p-value < 0.001 (***), p-value < 0.0001 (****).

To investigate the long-term stability of the CB phenotype, single cell clones of both lineages were culture-expanded and the phenotype of the resulting clones was analyzed. CD8αα^+^ and DN clones were frequently observed in UTC-derived clones, whereas CTC-derived clones remained predominantly CD8αβ^+^ (fig. 7C-D). This confirms that downregulation of CD8(β), and therefore the expression of CD8αα homodimers, is an exclusive characteristic of UTCs. Although Helios was induced in CTC-derived clones, expression remained significantly higher in UTC-derived clones. Similarly, CD26 expression was induced on UTC-derived clones, but remained significantly higher in CTC-derived clones (fig. 7E). In conclusion, the markers CD26, Helios, CD8α, CD8β and KIR reliably discriminated between UTCs and CTCs, even after culture expansion and activation.

Functional testing of the CD3^+^ TCRγδ^-^ CCR7^-^ CD26^-^ population revealed *ex vivo* CD3-induced killing activity. Upon activation, a spectrum of chemokines including IL-8, MIP-1α (CCL3), MIP-1β (CCL4) and fractalkine (CX3CL1) and the cytokines IL-2, FLT-3L, PDGF-AA, GM-CSF, IL-10, IFN-γ and TNFα was produced (fig. S5F-H).

## DISCUSSION

In the present study, the peripheral progeny of the human thymic IELps was identified in CB as a separate population evidently distinct from conventional CD8^+^ T cells and from NKT/MAIT cells. This population was remarkably heterogeneous. Five clusters were identified of which the MME^+^ UTCs and the GNLY^-^ MME^-^ UTCs are the closest related progeny of the thymic CD10^+^ PD-1^+^ population and are themselves the precursor clusters of the three differentiated clusters. It is therefore possible that in adulthood, when the output of the thymus diminishes, these two precursor populations gradually differentiate into effector cells and are no longer detectable in the blood and tissues. Of the effector clusters, the GNLY^+^ UTC and IL32^+^ UTC both contain cells expressing NK receptors including KIRs. KIR^+^ cells were not detected in the CTC clusters, *ex vivo* or after culture, indicating that KIR expression is confined to the uIEL lineage cells. A fifth UTC cluster, adjacent to the CTCs, was notable for high *GZMK* expression and low expression of *CD8A* and *CB8B*. A distinctive set of markers to further study this cluster was not found. Although some characteristics of the CTCs were present, it was argued that the GZMK^+^ DN UTC cluster also belongs to the uIEL lineage, because GZMK^+^ DN UTCs consistently co-occurred with the UTC population either in the CD3^+/low^ PD-1^+^ or CCR7^-^ EVI2B^+^ sorts and constitutively expressed low levels of CD8α and CD8β, in contrast to CTCs. Moreover, CD8^+^ UTCs have the discriminating ability to downregulate or lose CD8β (and CD8α) expression after *in vitro* stimulation whereas this was not observed for CD8^+^ CTCs.

The uIEL lineage in CB was likewise heterogeneous with regard to the expression of hallmark protein markers such as CD8β, PD-1, Helios and NK receptors. Despite the variable phenotypes within the uIEL population, evidence is presented that this population is well defined in CB with the markers CD26 and CCR7: the CCR7^-^ fraction of TCRαβ^+^ CD8^+^ cells consisted exclusively of CD26^-^ uIELs and CD26^+^ NKT/MAIT cells. However, CCR7 and to a lesser extent CD26 and Helios expression was unstable during T cell activation, which made these markers less valuable for analysis of samples containing antigen-experienced T cells such as adult blood. Hence, the presence of KIR^+^ T cells, CD8αα^+^ cells and DN T cells (excluding TCRγδ and NKT/MAIT cells), other markers that defined subtypes of this lineage and were not affected by T cell activation, was investigated and validated as a characteristic of the uIELs.

It was evidenced in CB that all KIR^+^ CD8^+^ T cells belong to the uIEL lineage, but only constitute a small fraction of the total uIEL population. Moreover, as CD8αα^+^ and DN cells could only be generated from UTCs and not from CTCs, KIR^-^ CD8αα^+^ and possibly DN T cells (after exclusion of TCRγδ^+^, NKT and MAIT cells) most likely also belong to the uIEL lineage. In addition to their transcriptional profile, it is therefore suggested that previously described populations, such as immunosuppressive KIR^+^ CD8^+^ T cells, KIR/NKG2A^+^ CD8^+^ T_VM_ cells and CD8αα^+^ cells described in tissues, probably belong to the unconventional, broadly reactive uIEL lineage rather than to the foreign peptide-reactive conventional CD8^+^ T cell lineage ^32–35,37^.

KIR^+^ CD8^+^ T cells were recently isolated from human adult blood based on their promiscuous binding to tetramers of HLA-A3 and HLA-A11. These cells express KLRC2, KIR2DL3 and NCR3, all of which are expressed by cells within the GNLY^*+*^ UTC and IL32^*+*^ UTC clusters. These cells are characterized by prominent expression of *IKZF2* and *ZNF683*, two characteristics of the uIEL lineage ^38^. KIR^+^ CD8^+^ cells, expressing *IKZF2*, are found to be enriched at inflammatory sites induced by viral infection as well as autoimmune inflammation in patients with celiac disease, multiple sclerosis and lupus. Importantly, these KIR^+^ CD8^+^ cells are shown to suppress autoreactivity by direct killing of pathogenic CD4 T cells ^39^. Functionally and phenotypically similar cells were described earlier in mice as CD8^+^ T_reg_ cells ^40–42^. Here, the uIELs were found to be able to kill target cells *ex vivo*. Furthermore, uIELs could produce chemokines (i.e. CCL3, CCL4 and CCL5), demonstrated as a mechanism to attract T cells ^43^. It is therefore possible that uIELs described here, may display *in vivo* immune suppressive properties by killing of autologous, metabolically active immune blasts. Indeed, it was previously reported that IELps are generated in the thymus by agonist selection at the early active DP blasts stage ^12^. As the IELps lineage is agonist selected on hematopoietic cells and not on thymic epithelial cells, it would be possible that IELps specifically react to autoantigens expressed by potent, metabolically active immune blasts ^44,45^. Therefore, whereas natural T_reg_ cells are reactive to antigens presented in the thymic medulla, the spectrum of antigens to which the uIELs react is probably presented by early DP blasts in the thymic cortex.

CD8αα^+^ cells have been described in various organs of the human body, including liver and lungs, and at tumor sites ^9,10,46,47^. CD8αα^+^ herpes simplex virus-specific T cells are found at the dermal-epidermal interface ^48^.

DN T cells represent a yet poorly characterized subset of TCRαβ^+^ T cells, partially due to their relatively low frequencies in human peripheral blood. They have mainly been studied in the context of autoimmunity, where increases of a heterogeneous group of DN T cells have been reported in patients with systemic lupus erythematosus and other autoimmune diseases. It is shown in human that peripheral DN T cells are maintained primarily by differentiation from CD8^+^ T cells ^49^. It is also reported in mice that only CD8^+^ T cells expressing PD-1 and Helios convert to DN T cells after encountering autoantigens ^37^. ScRNA-seq of splenic DN T cells in mice revealed five DN clusters of which two clusters were characterized by high expression of *IKZF2, IL2RB*, NK receptor genes such as *KLRD1* and chemokines such as *XCL1* and *CCL5*, genes also overexpressed in the uIEL lineage cells described here ^50^. Despite their low frequencies, DN T cells are potent producers of cytokines and therefore essential for immune responses. The therapeutic potential of DN T cells as Chimeric antigen receptor (CAR)-T cell is recently explored. It is evidenced that DN CAR-T cells are as effective as conventional CAR-T cells, without inducing toxicity, demonstrating the potential of using allogeneic DN T cells ^51^. A relationship between these DN T cells and the uIELs has not yet been conclusively addressed.

Finally, T_VM_ cells are described in mice as a subtype of CCR7^+^ conventional T cells with somewhat higher affinity for self-ligands in the thymus. The murine T_VM_ phenotype is imposed by self-reactivity of the TCR, similar to IELps. These cells start to express Eomes upon leaving the thymus and become activated in the periphery, proliferate on IL-15 and home to the tissues ^32–34^. The human counterpart is reported to be KIR/NKG2A^+^ Eomes^+^ and was detected in adult blood as well as CB samples ^35^. Here, it is reported that indeed a population is present in the UTC clusters in CB that weakly expresses Eomes and either KIRs or NKG2A.

In conclusion, the full spectrum of the uIEL lineage present in human CB is here described. The more immature uIELs, derived from thymic IELps, differentiate further and generate a heterogeneous mixture of effector cells. This population includes excellent killers, which may contribute to immune defence activity as well as exert immune suppressive activity by killing autologous immune blasts. The concept of these cells originating in the thymus by agonist selection as a consequence of the special characteristics of their TCR, may throw a different light on the study of these cells in immune reactivity to pathogens as well as autoimmunity.

## MATERIALS AND METHODS

### Sample Processing

Human CB was obtained from the Cord Blood Bank UZ Gent. Samples were used following the guidelines of the Medical Ethical Committee of Ghent University Hospital (CG20171208A, 8 December 2017) after informed consent had been obtained in accordance with the Declaration of Helsinki. Mononuclear cells were isolated using density gradient centrifugation (LymphoPrep; Axis-Shield, 1114547) and were enriched by magnetically activated cell sorting (MACS) through negative selection using anti-CD4-biotin, anti-CD14-biotin, anti-CD19-biotin, anti-CD235-biotin (homemade) and anti-biotin Microbeads (Miltenyi Biotec, 130-090-485). Human postnatal thymus was processed as previously described ^36^.

### Flow Cytometry and Antibodies

Staining of surface markers was performed in DPBS (Lonza, 17-512F) with 1% fetal calf serum (FCS; Biowest, S1810) using the antibody to cell ratio recommended by the supplier. Intracellular and intranuclear stainings were performed following the supplier’s protocol using BD Cytofix&Cytoperm (BD Biosciences, 554714) and the eBioscience™ Foxp3 / Transcription Factor Staining Buffer Set (eBioscience, 00-5523-00) respectively. Flow cytometric analysis was performed on the LSR II and cell sorting on the FacsARIA Fusion (both BD Biosciences). Flow cytometry data were analyzed using FACS DIVA software (BD Biosciences) and FlowJo software (TreeStar Inc). Viable cells were gated based on propidium iodide (PI) negativity or Fixable Viability Dye (eFluor 506; Thermo Fisher Scientific, 65-0866-18) negativity for surface and intracellular stainings respectively. The following list of anti-human monoclonal antibodies was used. Allophycocyanin (APC)/AF647-conjugated: CD4 (Miltenyi, 130-113-250), CD158b (KIR2DL2/DL3, Miltenyi, 130-092-617), Helios (Biolegend, 137221), Granzyme K (Biolegend, 370503), NKG2A (CD159a, Miltenyi, 130-114-089), PD-1 (CD279, Biolegend, 367420), TCRγδ (Miltenyi, 130-113-500); APC Cy7/APC Fire750-conjugated: CD8α (Biolegend, 344746), CCR7 (CD197, Biolegend, 353246); Brilliant Violet 421-conjugated: CD3 (Biolegend, 317344), CD54 (Biolegend, 353131); Brilliant Violet 605-conjugated: CD161 (Biolegend, 339915); Brilliant Violet 650-conjugated: CD3 (Biolegend, 317323); Brilliant Violet 711-conjugated: CCR7 (CD197, Biolegend, 353227); Brilliant Violet 785-conjugated: CD10 (Biolegend, 312237); Fluorescein isothiocyanate (FITC)-conjugated: CD8α (homemade), CD161 (Miltenyi, 130-114-118), IFN-γ (BD Biosciences, 554551), TCRγδ (BD Biosciences, 347903); Phycoerythrin (PE)-conjugated: CD158a/h (KIR2DL1/DS1, Miltenyi, 130-116-975), CD158b1/b2 (KIR2DL2/DL3, Beckman Coulter, IM2278U), CD158e1/e2 (KIR3DL/DS1, Beckman Coulter, IM3292), EVI2B (CD361, Thermo Fisher Scientific, A15806), Granzyme B (eBioscience, 12-8899-42), Granzyme K (Biolegend, 370511), PD-1 (CD279, Biolegend, 367404), Perforin (eBioscience, 12-9994-42); PE Cy7-conjugated: CD8β (eBioscience, 25-5273-42), CD10 (Biolegend, 312214), Eomes (eBioscience, 25-4877-41), TCRγδ (Biolegend, 331222); Peridinin chlorophyll protein complex (PerCP) Cy5.5-conjugated: CD4 (Biolegend, 344608), CD26 (Biolegend, 302715). CB MNCs were stained with Vδ1 (clone A13 supernatant, which can bind to Vδ1 when incorporated in hybrid Vδ1-Jα-Cα TCR chains, a kind gift from Prof. Dr. Lorenzo Moretta’s laboratory), anti–mouse Ig light chain κ (Biolegend, 409506), 5% normal mouse serum (Invitrogen, 10410), followed by the appropriate antibodies above to isolate the populations.

CD4/CD14/CD19/CD235-depleted CB MNCs were sorted into CD3^+^ TCRγδ^-^ CD4^-^ CD8α^+^ PD-1^-^ (PD-1^-^ population) and CD3^+/low^ TCRγδ^-^ CD4^-^ CD8α^+^ PD-1^+^ (PD-1^+^ population) for the transcriptome, TCR and proteome analyses. The PNT populations were sorted as previously described ^36^. For the single cell assay, CD4/CD14/CD19/CD235-depleted CB MNCs were sorted into CD3^+/low^ TCRγδ^-^ CD4^-^ CD8α^+^ PD-1^+^ (sort 1), CD3^+/low^ TCRγδ^-^ CD4^-^ CD8α^+^ CCR7^-^ EVI2B^+^ (sort 2), CD3^+^ TCRγδ^-^ CD4^-^ PD-1^-^ and CD3^+/low^ TCRγδ^-^ CD4^-^ PD-1^+^ (sort 3), CD3^+^ TCRγδ^-^ CD4^-^CCR7^+^ EVI2B^-^ and CD3^+/low^ TCRγδ^-^ CD4^-^ CCR7^-^ EVI2B^+^ (sort 4). For the proliferation and stimulation experiments, CD4/CD14/CD19/CD235-depleted CB MNCs were sorted into CD3^+^ TCRγδ^-^ CD4^-^ CD8α^+^ CCR7^+^ (CTC) and CD3^+/low^ TCRγδ^-^ CD4^-^ CCR7^-^ CD26^-^ (UTC).

### RNA Sequencing

The populations of interest were each time sorted from three CB and PNT donors. The CD3^+^ TCRγδ^+^ population was also sorted, only 2 TCRγδ^+^ samples were analyzed for CB. The PNT and CB populations were sorted in IMDM (Thermo Fisher Scientific, 12440053) supplemented with 10% FCS, 2 mM L-glutamine (Thermo Fisher Scientific, 25030-081), 100 IU/mL penicillin and 100 IU/mL streptomycin (Thermo Fisher Scientific, 15140-122) (complete IMDM, cIMDM) and washed 3 times in phosphate-buffered saline (PBS). RNA extraction was performed using the miRNeasy Mini Kit (Qiagen, 217004). For poly(A) RNA-seq, the QuantSeq 3′ mRNA FWD kit (Lexogen) was used, followed by single-ended sequencing on the NextSeq500 Sequencing System (Illumina) with a read length of 75bp. RNA-seq reads were aligned to hg38-noalt using STAR v2.6.0c and quantified on Ensembl v93.

### TCR sequencing

For the TCR analysis, the populations of interest were sorted from six CB donors and two new PNT donors. Previously published PNT samples were also reanalyzed (12). RNA extraction was performed using the miRNeasy Micro Kit (Qiagen, 217084), followed by template-switch anchored RT-PCR. High-throughput sequencing of TRA and TRB loci was performed as previously described ^52^. Raw sequencing reads from fastq files were aligned to reference V, D and J genes from GenBank database specifically for ‘TRA’ or ‘TRB’ to build CDR3 sequences using the MiXCR software version 3.0.12 ^53^. Following, the CDR3 sequences were analysed using VDJtools software version 1.2.1 ^54^. Out of frame sequences were excluded from the analysis, as well as non-functional TRA and TRB segments using *IMGT* (the international ImMunoGeneTics information system®) annotation. TRDV gene segment-containing sequences were filtered as well, except for the analysis where the amount of TRDV1 containing sequences was assessed. Calcbasicstats default function was used to calculate the number of CDR3 N additions. Cumulative gene segment plots were generated using the output from CalcSegmentUsage function. Tree maps were generated using the Treemap Package on RStudio, grouping TRAV and TRAJ segments according to their locus position. D75 repertoire diversity metrics were calculated by measuring the percentage of clonotypes required to occupy 75% of the total TCR repertoire. Determination of the CDR3α and CDR3β apex region and cysteine usage was performed following previously described indications ^14^. Hydrophobic CDR3α and CDR3β doublet containing sequences were determined by calculating the percentage of sequences using any of the 175 amino acid doublets previously identified as promoting self-reactivity ^13^. Physicochemical characteristics (strength, volume and polarity) of the CDR3β were analysed using VDJtools software version 1.2.

### LC-MS/MS proteomic analysis

#### Sample preparation

The populations of interest were each time sorted from three CB and PNT donors. Cell pellets (± 1.10^6^ cells per pellet) were resuspended in lysis buffer (8 M urea; 20 mM HEPES, pH 8.0). Samples were sonicated by three pulses of 15 s, interspaced by 1 min pauses on ice at an intensity output of 15 W, and centrifuged for 15 min at 20 000g at room temperature to remove insoluble components. Proteins were reduced with 5 mM dithiothreitol (DTT) (Sigma-Aldrich) for 30 min at 55 °C and then alkylated by the addition of 10 mM iodoacetamide (Sigma-Aldrich) for 15 min at room temperature in the dark. Samples were further diluted with 20 mM HEPES, pH 8.0, to a final urea concentration of 4 M and proteins were digested with LysC (Wako) (1/100, w/w) for 4 hours at 37°C. Samples were again diluted to 2 M urea and digested with trypsin (Promega) (1/100, w/w) overnight at 37°C. The resulting peptide mixture was acidified by addition of 1% trifluoroacetic acid (TFA). Peptides were then purified on a SampliQ SPE C18 cartridge (Agilent), vacuum-dried, and kept at −20 °C until measured by LC-MS/MS.

#### LC–MS/MS Analysis

Immediately before injection, purified peptides were redissolved in 15 μL loading solvent (0.1% trifluoroacetic acid/water/acetonitrile (0.1:98:2, v/v/v)) and the peptide concentration was determined by measuring on a Lunatic spectrophotometer (Unchained Labs). 2 μg of peptide material of each sample was injected for LC-MS/MS analysis on an Ultimate 3000 RSLC nano-LC (Thermo Fisher Scientific, Bremen, Germany) in-line connected to a Q Exactive HF mass spectrometer (Thermo Fisher Scientific) equipped with a nanospray flex ion source (Thermo Fisher Scientific). Trapping was performed at 10 μL/min for 4 min in loading solvent A on a 20-mm trapping column (made in-house, 100-μm internal diameter, 5-μm beads, C18 Reprosil-HD, Dr Maisch, Germany). Peptide separation after trapping was performed on a 200-cm-long micropillar array column (PharmaFluidics) with C18-endcapped functionality. The Ultimate 3000’s column oven was set to 50°C. For proper ionization, a fused silica PicoTip emitter (10-μm inner diameter) (New Objective) was connected to the μPAC outlet union and a grounded connection was provided to this union. Peptides were eluted by a nonlinear gradient from 1 to 55% MS solvent B (0.1% FA in water/acetonitrile (2:8, v/v)) over 145 min, starting at a flow rate of 750 nL/min switching to 300 nL/min after 15 min, followed by a 15-min washing phase plateauing at 99% MS solvent B. Re-equilibration with 99% MS solvent A (0.1% FA in water) was performed at 300 nL/min for 45 min followed by 5 min at 750 nL/min adding up to a total run length of 210 min. The mass spectrometer was operated in a data-dependent, positive ionization mode, automatically switching between MS and MS/MS acquisition for the 16 most abundant peaks in each MS spectrum. The source voltage was 2.2 kV, and the capillary temperature was 275 °C. One MS1 scan (m/z 375–1,500, AGC target 3.10^6^ ions, maximum ion injection time 60 ms), acquired at a resolution of 60,000 (at 200 m/z), was followed by up to 16 tandem MS scans (resolution 15,000 at 200 m/z) of the most intense ions fulfilling predefined selection criteria (AGC target 1.10^5^ ions, maximum ion injection time 80 ms, isolation window 1.5 Da, fixed first mass 145 m/z, spectrum data type: centroid, intensity threshold 1.3 × 10^4^, exclusion of unassigned, 1, 7, 8, >8 positively charged precursors, peptide match preferred, exclude isotopes on, dynamic exclusion time 12 s). The higher-energy collisional dissociation was set to 28% normalized collision energy, and the polydimethylcyclosiloxane background ion at 445.12003 Da was used for internal calibration (lock mass).

#### Data analysis

Data analysis was performed with MaxQuant (version 1.6.2.6) using the Andromeda search engine with default search settings including a false discovery rate (FDR) set at 1% on both the peptide and protein level. Spectra were searched against the human Swiss-Prot database (from November 2018 with 20 424 entries) separately for PNT and CB T cells. The mass tolerance for precursor and fragment ions was set to 4.5 and 20 ppm, respectively, during the main search. Enzyme specificity was set to the C-terminal of arginine and lysine, also allowing cleavage next to prolines with a maximum of two missed cleavages. Variable modifications were set to oxidation of methionine residues and acetylation of protein N-termini. Matching between runs was enabled with a matching time window of 1.5 min and an alignment time window of 20 min. Only proteins with at least one unique or razor peptide were retained, leading to the identification of 4539 and 3584 proteins for PNT and CB, respectively. Proteins were quantified by the MaxLFQ algorithm integrated into the MaxQuant software. A minimum ratio count of two unique or razor peptides was required for quantification. Further data analysis was performed with the Perseus software (version 1.6.2.1) separately for the PNT and CB data set after uploading the protein groups file from MaxQuant. Reverse database hits, potential contaminants and proteins that are only identified by peptides carrying at least one modified amino acid were removed. Replicate samples were grouped and proteins with less than three valid values in at least one group were removed, and missing values were imputed from a normal distribution around the detection limit resulting in 3,001 and 2,024 quantified proteins for PNT and CB, respectively.

### Gene set enrichment analysis

GSEA was performed using the GSEA software version 4.1.0., a joint project of UC San Diego (San Diego, CA, USA) and Broad Institute (Cambridge, MA, USA)^55,56^. The GSEAPreranked tool was run using standard parameters and 1000 permutations. The gene set contained the significantly differentially expressed genes when comparing the CD10^+^ PD-1^+^ IELp population to the CD10^−^ PD-1^-^ cells from human PNT. The gene list was ranked by comparing the PD-1^+^ population to the PD-1^-^ cells from human CB, ranked from the upregulated genes (left) to the downregulated genes (right). The normalized enrichment score (NES) reflects the degree to which the gene set is overrepresented in the upregulated genes (positive value) or downregulated genes (negative value). The false discovery rate q value (FDR q) is the estimated probability that a gene set with a given NES represents a false-positive finding.

### Single-cell RNA sequencing analysis

#### Single cell library preparation and sequencing

The populations of interest were sorted from CD4/CD14/CD19/CD235-depleted CB from two different donors. From the first donor, CD3^+/low^ TCRγδ^-^ CD4^-^ CD8α^+^ as CD3^+/low^ PD-1^+^ (sort 1) or CCR7^-^ EVI2B^+^ (sort 2) were sorted separately. Both fractions were labeled with different TotalSeq anti-human Hashtag antibodies (Biolegend) before being pooled in equal portions and processed together. From the second donor, CD3^+^ TCRγδ^-^ CD4^-^ PD-1^-^ and CD3^+/low^ TCRγδ^-^ CD4^-^ PD-1^+^ (sort 3), CD3^+^ TCRγδ^-^ CD4^-^ CCR7^+^ EVI2B^-^and CD3^+/low^ TCRγδ^-^ CD4^-^ CCR7^-^ EVI2B^+^ (sort 4) were sorted. Considering the smaller percentage of unconventional T cells, the conventional cells were sorted separately and later added in equal portions. Sort 4 was combined with CITE-seq labeling. Cells were incubated for 30 min on ice with 50 μL of staining mix in PBS containing 0.04% BSA, Fc receptor block (PN 422301, TruStain FcX, BioLegend) and a human cell surface protein antibody panel containing 277 oligo-conjugated antibodies (TotalSeq-A, BioLegend) including 6 TotalSeq-A isotype controls (table S4). TotalSeq antibodies were diluted in concentrations as recommended by the manufacturer. Sorted single-cell suspensions were resuspended at an estimated final concentration of 1200 cells/μl and loaded on a Chromium GemCode Single Cell Instrument (10x Genomics) to generate single-cell gel beads-in-emulsion (GEM) at the VIB Single Cell Core.The scRNA-Seq libraries were prepared using the GemCode Single Cell 3’ Gel Bead and Library kit, version NextGEM 3.1 (10x Genomics) according to the manufacturer’s instructions with the addition of amplification primers (3nM), 5’CCTTGGCACCCGAGAATT*C*C and 5’GTGACTGGAGTTCAGACGTGTGC*T*C during cDNA amplification to enrich the TotalSeq-A cell surface and hashtag protein oligos. Library construction was performed according to the manufacturer’s instructions. Sequencing libraries were loaded on an Illumina NovaSeq flow cell at VIB Nucleomics core with sequencing settings according to the recommendations of 10x Genomics, pooled in a 80:25 ratio for the combined 3’ gene expression and cell surface protein libraries, respectively.

#### Preprocessing of the scRNA-seq and CITE-seq data

The Cell Ranger pipeline (10x Genomics, version 3.1.0) was used to perform sample demultiplexing and to generate FASTQ files for read 1, read 2 and the i7 sample index for the gene expression and cell surface protein libraries. Read 2 of the gene expression libraries was mapped to the reference genome (GRCh38.99) using STAR. The resulting count matrices were subsequently loaded into R for further processing using Seurat version 4.0.5 ^57^. Empty and/or damaged cells were removed from the datasets by filtering on the following three parameters: (i) number of genes per cell (nFeature > 600), (ii) number of UMI counts per cell (nCount > 1150) and (iii) percentage mitochondrial genes per cell (percent.mt < 15). The remaining cells were normalized with the built-in normalization function from the Seurat package using the log normalization method and a scale factor of 10000. Highly variable features/genes were identified with FindVariableFeatures with the selection method set to vst and the nfeatures parameter to 4000. Differential gene expression analysis between cell clusters and conditions was performed using Wilcoxon Rank Sum test through the Seurat function “FindMarkers”. P-value adjustment was performed using Bonferroni correction.

CITE-seq antibody reads were quantified using the feature-barcoding functionality within the Seurat package. Antibodies with low expression were filtered out based on inspection of the feature plots for each antibody. After processing, the CITE-Seq and scRNA-seq data of Sort 4 were merged into the same Seurat object.

#### Dataset integration and batch correction

To verify that our different sorting strategies/definitions of the uIEL in CB included the same or different cell types, a data integration of our three CB samples was performed using the Seurat package. Therefore, the integration anchor strategy was followed. Integration anchors were identified following the reciprocal PCA (rPCA) method instead of the CCA method, due to the speed of the former method and the recommendation of the developers that rPCA is more conservative and thus better equipped to handle cell populations that have no matching type between samples. For this integration, integration features were first selected with the “SelectIntegrationFeatures” function and subsequently the samples were scaled (“ScaleData”) and a PCA analysis (“RunPCA”) was executed for each sample separately. Afterwards the integration anchors were identified with the “FindIntegrationAnchors” function. After integration with the “IntegrateData” function the data was scaled once again and a new PCA analysis was performed for visualization and data exploration.

Subsequently, to identify the progeny of the thymic uIEL lineage in CB a second data integration was performed with CB and PNT samples. Therefore, four PNT samples (TTA9, TTA10, TTA12 and TTA14) were selected from Park et al. ^15^. The FASTQ-files of these four samples were acquired and processed them as described under ‘Preprocessing of the scRNA-seq and CITE-seq data’. By processing the samples of both organs in the same way, the variation that needed to be corrected in the subsequent data integration and batch correction step, could be limited. For the latter, the R package Harmony was used ^58^. Prior to the data integration with Harmony all samples were merged into a combined Seurat object afterwards the following steps were performed: (i) identification of highly variable genes/features (“FindVariableFeatures”), (ii) data scaling (“ScaleData”) and (iii) PCA analysis (“RunPCA”). The data integration was executed with the function “RunHarmony”, taking into account three sources of potential batch effects, namely donor, sort and chemistry. The later was added since the PNT samples were processed with the v2 version of the 10X genomics scRNA-seq kit, while for the CB samples the v3 version was used. Afterwards, the integrated object was processed for visualization and data exploration.

#### Trajectory Analysis and SCENIC

Based on the data integration of the PNT and CB samples, a trajectory analysis could be performed to identify the progeny of the uIELs. For this trajectory analysis, the TSCAN algorithm (version 1.28.0) was used ^59^. To simplify the problem for the algorithm, the populations of interest were selected prior to the actual analysis. For the unconventional populations, these selected cell types were: CD8αα(II), MME^+^ UTC, GNLY^-^ MME^-^ UTC, GNLY^+^ UTC, GZMK^+^ DN UTC, and IL32^+^ UTC. While for the conventional populations these were: CD8^+^T, CTC1, CTC2, CTC3, CTC4, and CTC5. The first step in the TSCAN analysis was the calculation of the centroid of each cluster with the “reducedDim” function. These centroids were then connected with a minimum spanning tree (MST) in the subsequent step using the “createClusterMST” function. The cells were subsequently mapped and ordered along this MST to determine their pseudo-time point. This was done with the functions “mapCellsToEdges” and “orderCells”. To identify the genes that might play a role in this differentiation process along the pseudo-time, a differential expression analysis with the tradeSeq package (version 1.4.0) was performed ^60^. Therefore, a generalized additive model (GAM) was fitted to the data along the pseudotime and subsequently the associationTest was performed to identify the differentially expressed genes.

To better identify which transcription factors are important or active in the different cell populations, the data was analyzed with the SCENIC algorithm ^61^. Note that for the actual analysis the python implementation of this package, pySCENIC, was used. In a first step gene regulatory networks and co-expression modules are generated by the GRNBoost 2 algorithm based on the correlation between transcription factors and other genes. Subsequently regulons are predicted based on known binding motifs provided by the Aerts lab. And in the final step the cellular enrichment for the different regulons was calculated using the Aucell algorithm. The output of the different steps was loaded into python to calculate the regulon specificity scores (RSS) for the different cell populations through the pySCENIC package.

The regulon specificity score (RSS) was calculated from the regulon activity score (RAS) and lies between 0 and 1. With a higher value for RSS indicating a higher specificity of that regulon in the cell type compared to others. The UTC and CTC specific regulons were selected according to the following strategy: a regulon that appeared in the top 5 of one of the UTC or CTC populations and in the top 10 of one or more of the remaining UTC or CTC populations was selected for inclusion in the plot. The RAS was calculated based on the rank of the expression value in the cell of all genes involved in the regulon ^62^.

### T cell expansion

The CellTrace proliferation assays were performed as previously described ^36^. CTC and UTC clones were generated by FACS sorting single cells and expanding them on irradiated allogenic feeder cells, consisting of a mixture of 40 Gy irradiated peripheral blood mononuclear cells and 50 Gy irradiated JY cells. Cells were cultured in cIMDM, supplemented with 1 μg/mL phytohemagglutinin (PHA, Sigma–Aldrich). IL-2 (5 ng/mL; Miltenyi, 130-097-748) was added on day five and day ten. Cells were restimulated every 7 to 14 days. After 14-28 days, grown clones were harvested and assessed via flow cytometry.

### ^51^Chromium Release Assay

Target cells (W6/32 or OKT3 hybridoma) were labelled with ^51^Chromium (Perkin Elmer) for 90 min at 37 °C, washed and added at 10^3^ cells per well to various ratios of effector T cells (CTC or UTC population after overnight incubation with IL-15) in 96 well V-bottomed plates (NUNC, Thermo Fisher Scientific). After 4 hours of co-incubation, the supernatant was harvested and measured in a 1450 LSC & Luminescence Counter (Perkin Elmer). Specific lysis was calculated as follows: (experimental release–spontaneous release)/(maximal release–spontaneous release) × 100%.

### Cytokine Production

To explore the secreted cytokine profile of the CB populations, Luminex High Performance Assays (R&D Systems) were performed. The supernatant of both freshly sorted UTCs and UTC-derived clones after 24 hours of stimulation with phorbol myristate acetate (PMA, 1 ng/mL; Sigma-Aldrich, 16561-29-8) + ionomycin (0.5 μg/mL; Sigma-Aldrich, 56092-82-1) was assessed with the Luminex Performance Human XL Cytokine Magnetic Panel 44-plex Fixed Panel (Bio-Techne, LKTM014). Following, a custom mixed multiplex was used to determine the IFN-γ, granzyme B, GM-CSF, MIP-1α, MIP-1β and IL-2 concentrations in the supernatant of both freshly sorted CTCs and UTCs after 24 hours of stimulation with PMA + ionomycin. All assays were performed conform the manufacturer’s protocol and measured with the Bio-Plex 200 system (Biorad) and analysed with the Bio-Plex Manager software version 6.2. Levels below or above the detection level were set as the lower or upper detection level, respectively.

### Statistical Analysis

Statistical analyses were performed in Prism version 9.3.1. (GraphPad Software, San Diego, CA, USA), using statistical tests as indicated in figure legends. Results were considered statistically significant when the p-value was less than 0.05.

## Data availability

The mass spectrometry proteomics data have been deposited to the ProteomeXchange Consortium via the PRIDE ^63^ partner repository with the dataset identifier PXD033392. The sequencing data discussed in this publication have been deposited in NCBI’s Gene Expression Omnibus ^64^ and will be made accessible upon publication through GEO Series accession number GSE201811.

## Supplementary Material

Fig. S1. RNA and protein expression profile by the PNT CD10^+^ PD-1^+^ population.

Fig. S2. Distinctive TCR repertoire of the PNT CD10^+^ PD-1^+^ population.

Fig. S3. Defining the T cell clusters in CB.

Fig. S4. Phenotyping the unconventional populations in CB.

Fig. S5. UTC phenotype and functionality after proliferation with IL-15.

Table S1. Significantly differentially expressed between the PD-1^+^ and PD-1^-^ population, in both PNT and CB

Table S2. Cell number in each identified cluster

Table S3. Top 10 differentially expressed genes for each identified cluster

Table S4. CITE-seq cell surface protein antibody panel

Table S5. Luminex Performance Human XL Cytokine Magnetic Panel 44-plex Fixed Panel

## Author contributions

Conceptualization: L. Billiet, L. De Cock, B. Vandekerckhove; Data curation: L. Billiet, L. De Cock, B. Vandekerckhove; Formal Analysis: L. Billiet, L. De Cock, G. Sanchez Sanchez, R. L. Mayer, F. Van Nieuwerburgh; Funding acquisition: D. Vermijlen, F. Impens, B. Menten, B. Vandekerckhove; Investigation: L. Billiet, L. De Cock, G. Sanchez Sanchez, R. L. Mayer, G. Goetgeluk, N. Vandamme; Methodology: L. Billiet, L. De Cock, B. Vandekerckhove; Project administration: L. Billiet, B. Vandekerckhove, Software: L. De Cock, N. Vandamme, R. Seurinck, J. Roels, M. Lavaert: Supervision: G. Leclercq, T. Taghon, F. Impens, B. Menten, D. Vermijlen, B. Vandekerckhove; Validation: B. Vandekerckhove; Visualization: L. Billiet, L. De Cock, G. Sanchez Sanchez, R.L. Mayer; Writing – original draft: L. Billiet, B. Vandekerckhove; Writing – review & editing: L. De Cock, G. Sanchez Sanchez, R. L. Mayer, S. De Munter, M. Pille, J. Ingels, H. Jansen, K. Weening, E. Pascal, K. Raes, S. Bonte, T. Kerre, N. Vandamme, R. Seurinck, J. Roels, F. Van Nieuwerburgh, G. Leclercq, T. Taghon, F. Impens, B. Menten, D. Vermijlen.

## Acknowledgments

This work is funded by Research Foundation Flanders (Fonds voor Wetenschappelijk Onderzoek—FWO, grant 3F012519), Stichting tegen Kanker (STK, grant FAF-F/2016/756), GOA (grant 2021.0008), FNRS CDR (grant CDR_J.0225.20) and the TCR GSS is supported by Télévie-FNRS (grant 7.4586.19 and 7.6529.21). The authors would like to thank Dr. Conny Matthys of the Cord Blood Bank UZ Gent, Sergi Rodà Llordés for his help and advice to optimize data filtering for analysis of the CDR3 sequences, Gabriële Holtappels for her expertise in Luminex High Performance Assays, Juliette Roels for her tips and tricks regarding R and the UGent Core Flowcytometrie for their help with flow cytometry and cell sorting. We thank Hilde Cheroutre, Derk Amsen and Greet Verstichel for critically reading the manuscript and their helpful suggestions.

The authors declare no competing financial interests.

## Abbreviations

CB: cord blood
CTC: conventional T cell
DN: double negative
GSEA: gene set enrichment analysis
IEL: intraepithelial lymphocyte
IELp: intraepithelial lymphocyte precursor
KIR: killer Ig-like receptor
MAIT: mucosal-associated invariant T cell
NKT: natural killer T cell
PNT: postnatal thymus
SP: single positive
TCR: T cell receptor
TF: transcription factor
T_VM_: virtual memory T cell
uIEL: unconventional intraepithelial lymphocyte
UMAP: uniform manifold approximation and projection
UTC: unconventional T cell

