## Supplemental materials for "Single-cell profiling identifies a spectrum of human unconventional intraepithelial T lineage cells"

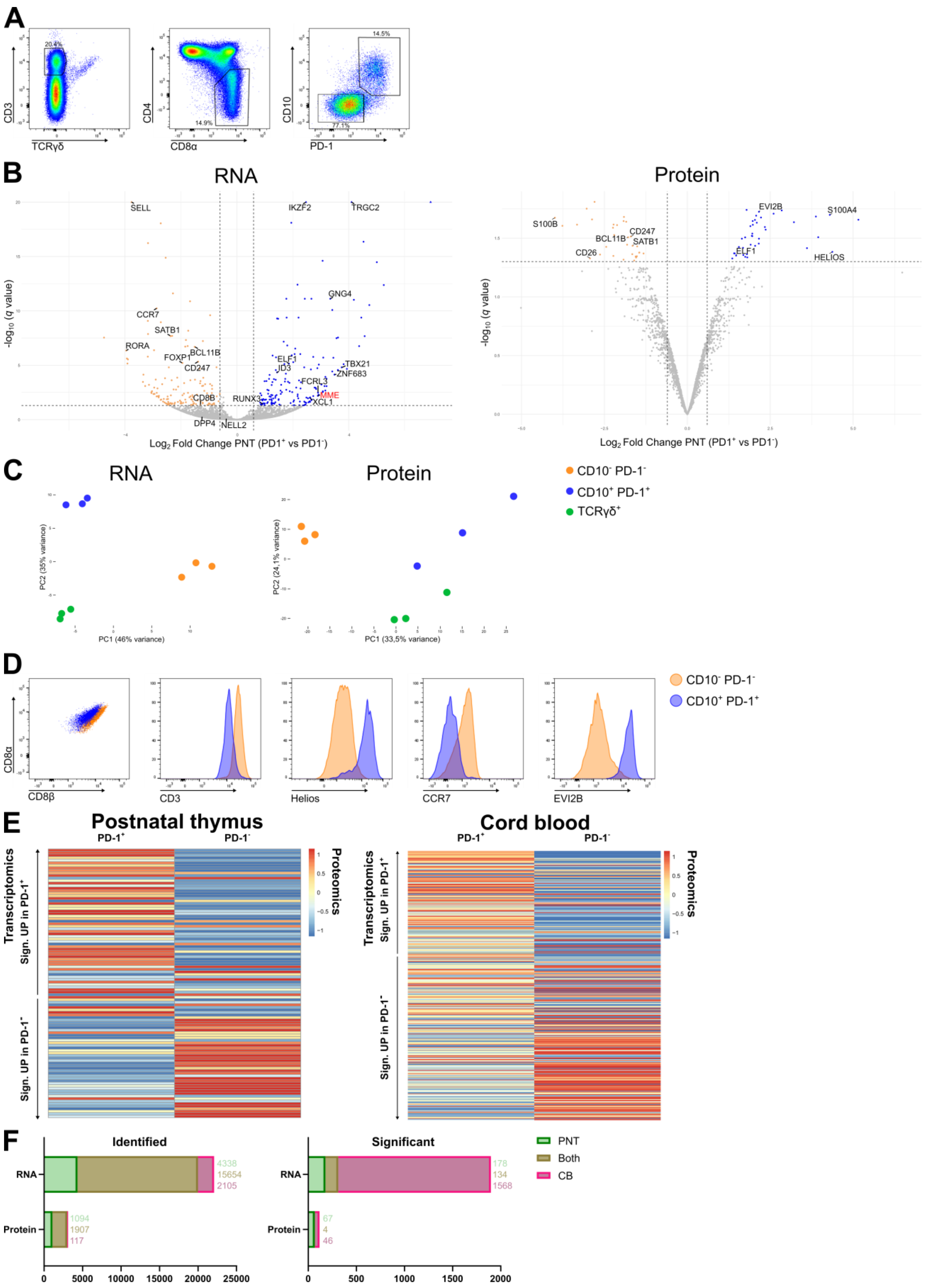

**Supplementary Fig. 1. RNA and protein expression profile by the PNT CD10$^+$ PD-1$^+$ population.** (**A**) Representative gating strategy for the CD3$^+$ TCRγδ$^-$ CD4$^-$ CD8α$^+$

CD10$^+$ PD-1$^+$ populations in human PNT. (**B**) Volcano plots of differentially expressed genes (left) and proteins (right) between the PNT CD10$^+$ PD-1$^+$ and CD10$^-$ PD-1$^-$ populations. Triangles indicate data points outside the y-axis range. Data points with a $|Log_2$ Fold Change$| > 0.6$ and adjusted $P < 0.05$ are colored (upregulated in blue, downregulated in orange). (**C**) PCA of the transcriptome (left, donor corrected) and proteome (right) analysis of the sorted populations from three different PNT donors. (**D**) Flow cytometric analysis of CD8α, CD8β, CD3, Helios, CCR7 and EVI2B on the CD10$^-$ PD-1$^-$ and CD10$^+$ PD-1$^+$ population in PNT, representative of at least three PNTs. (**E**) Heatmap showing the corresponding relative protein abundance of all significantly differentially expressed genes between the PNT CD10$^+$ PD-1$^+$ and CD10$^-$ PD-1$^-$ population (left) or between the CB CD3$^{+/low}$ PD-1$^+$ and CD3$^+$ PD-1$^-$ population (right). Differentially expressed genes of which the corresponding protein was not detected are not shown. The heatmap shows column- and row-scaled, normalized, mean protein abundance. (**F**) Bar graph showing the overlap in identified (left) and significantly differentially expressed (right) RNAs and proteins in PNT and CB populations.

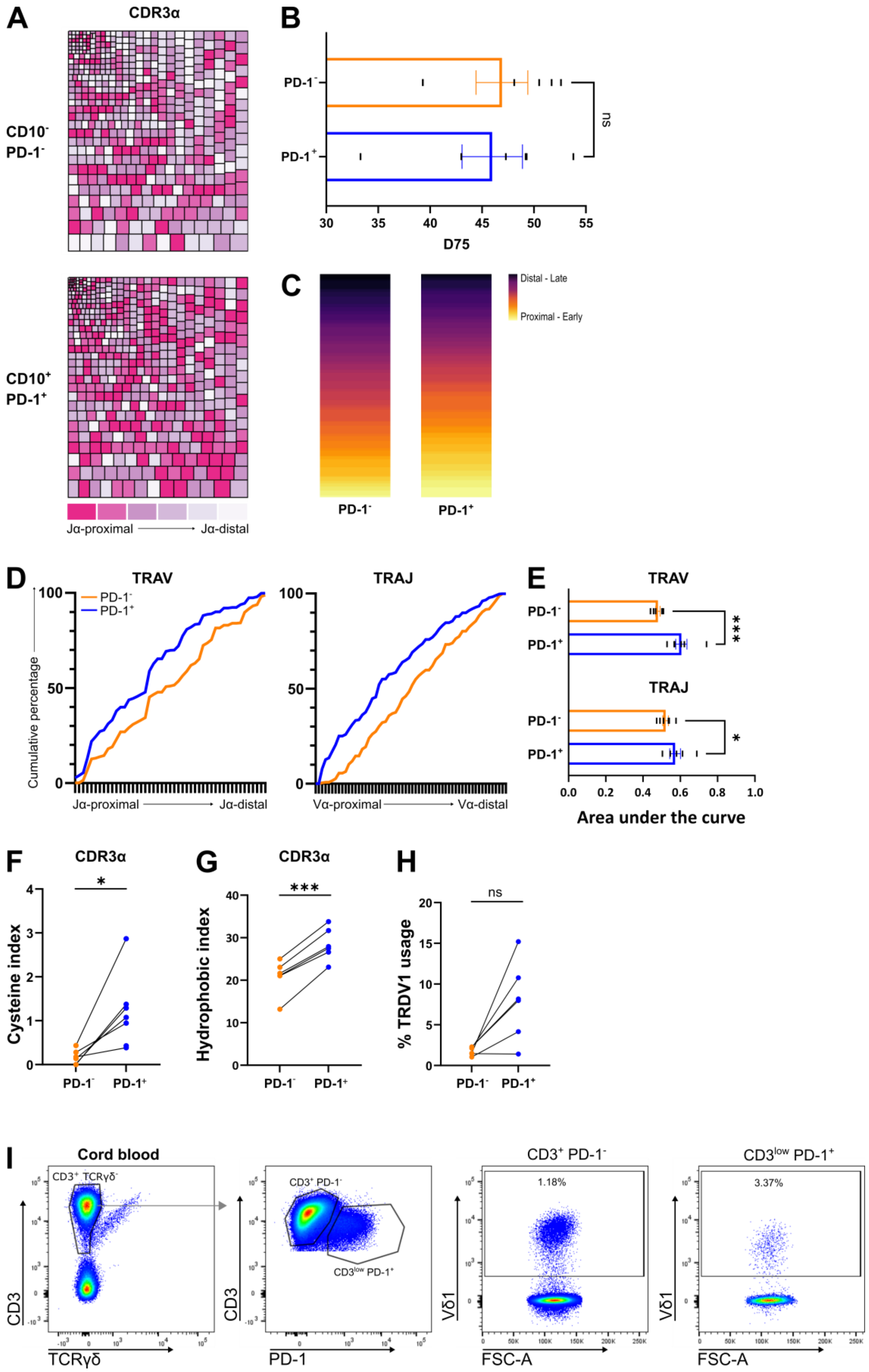

**Supplementary Fig. 2. Distinctive TCR repertoire of the PNT CD10$^+$ PD-1$^+$ population.** (**A**) Representative tree maps showing CDR3α clonotype usage in relation to repertoire size for the CD10$^-$ PD-1$^-$ (top) and CD10$^+$ PD-1$^+$ (bottom) populations. Each rectangle represents one CDR3 clonotype and its size corresponds to its relative frequency in the repertoire (rectangle colors are categorized from J-proximal (pink) to J-distal (white) for the TRAV gene segments). (**B**) D75 (percentage of clonotypes required to occupy 75% of the total TCR repertoire) analysis comparing the PD-1$^-$ and PD-1$^+$ population (individual values and mean ± SEM). Mean difference was not significant. (**C**) Representative heatmap illustrating the difference in TCR Vα usage between the PD-1$^-$ and PD-1$^+$ population. (**D**) Representative cumulative percentage of TRAV (left) or TRAJ (right) gene segment usage. X-axis represents the location in the TRAV or TRAJ locus. (**E**) Area under the curve determined from the cumulative plots from each sample (individual values and mean ± SEM). Šídák's multiple comparisons test was used to assess the statistically significant difference. p-value < 0.05 (*), p- value < 0.001 (***). (**F**) Cystein index (percentage of unique sequences with cysteine within 2 positions of the CDR3 apex) and (**G**) Hydrophobic index (percentage of unique sequences with self-reactive hydrophobic CDR3 position 6 and 7 doublets) of the CDR3α. (**H**) Percentage of unique sequences containing a *TRDV1* segment. (**B, F-H**) Paired t-tests were used to assess statistical significance. Connected values correspond to paired populations of the same biological replicate (n= 6). p-value > 0.05 (ns), p-value < 0.05 (*), p-value < 0.001 (***). (**I**) Representative flow cytometric analysis of the Vδ1$^+$ (A13 clone) cells in both CB populations.

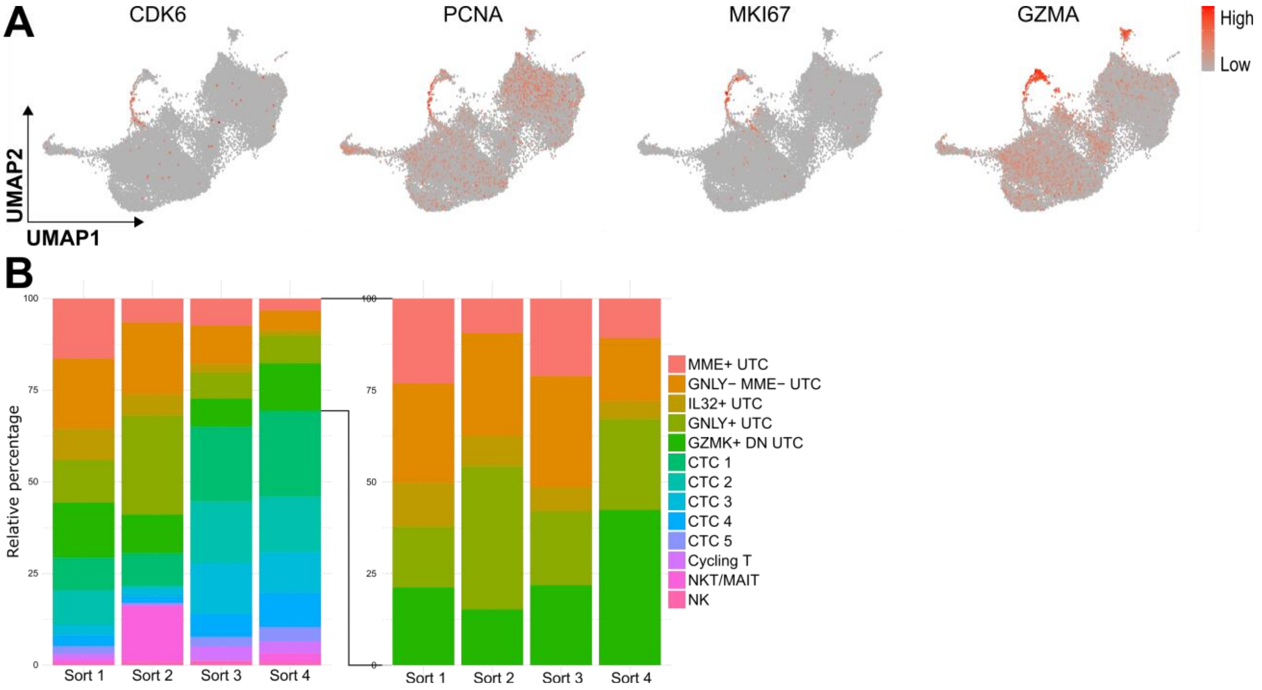

**Supplementary Fig. 3. Defining the T cell clusters in CB. (A)** UMAP feature plots of characteristic genes for the cycling T cells. (**B**) Stacked bar chart showing the composition of the 13 identified clusters (left) or the five different UTC clusters (right) in the different sorts.

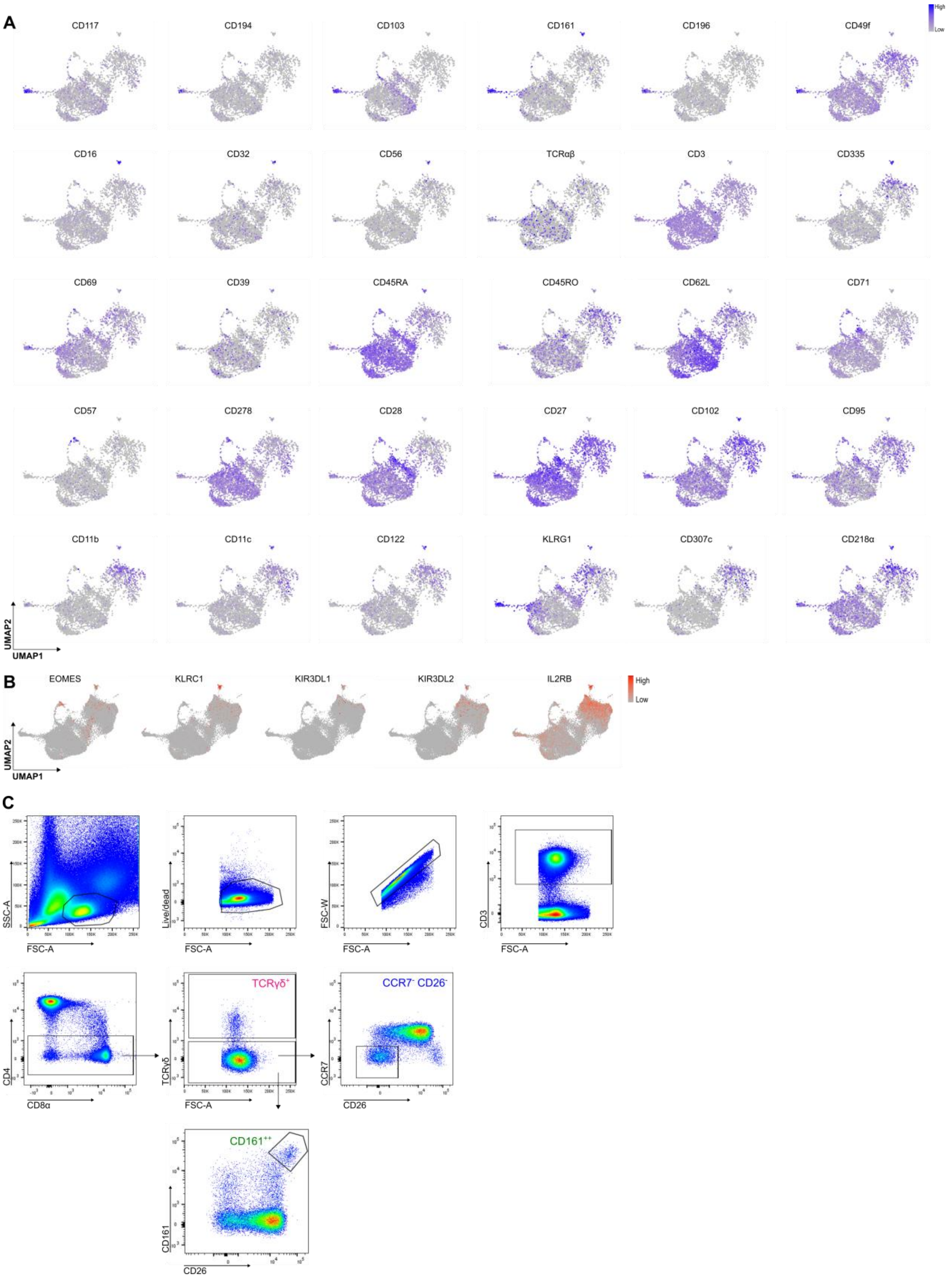

**Supplementary Fig. 4**. **Phenotyping the unconventional populations in CB.** (**A**) Protein-based UMAP visualizations showing the expression of the indicated cell surface protein markers, only visualizing the single cells from which protein data was collected. (**B**) UMAP feature plots of characteristic genes previously appointed to virtual memory T cells. (**C**) Flow cytometric gating strategy for the unconventional populations in CB. Gating strategy used to obtain the percentages in figure 6F.

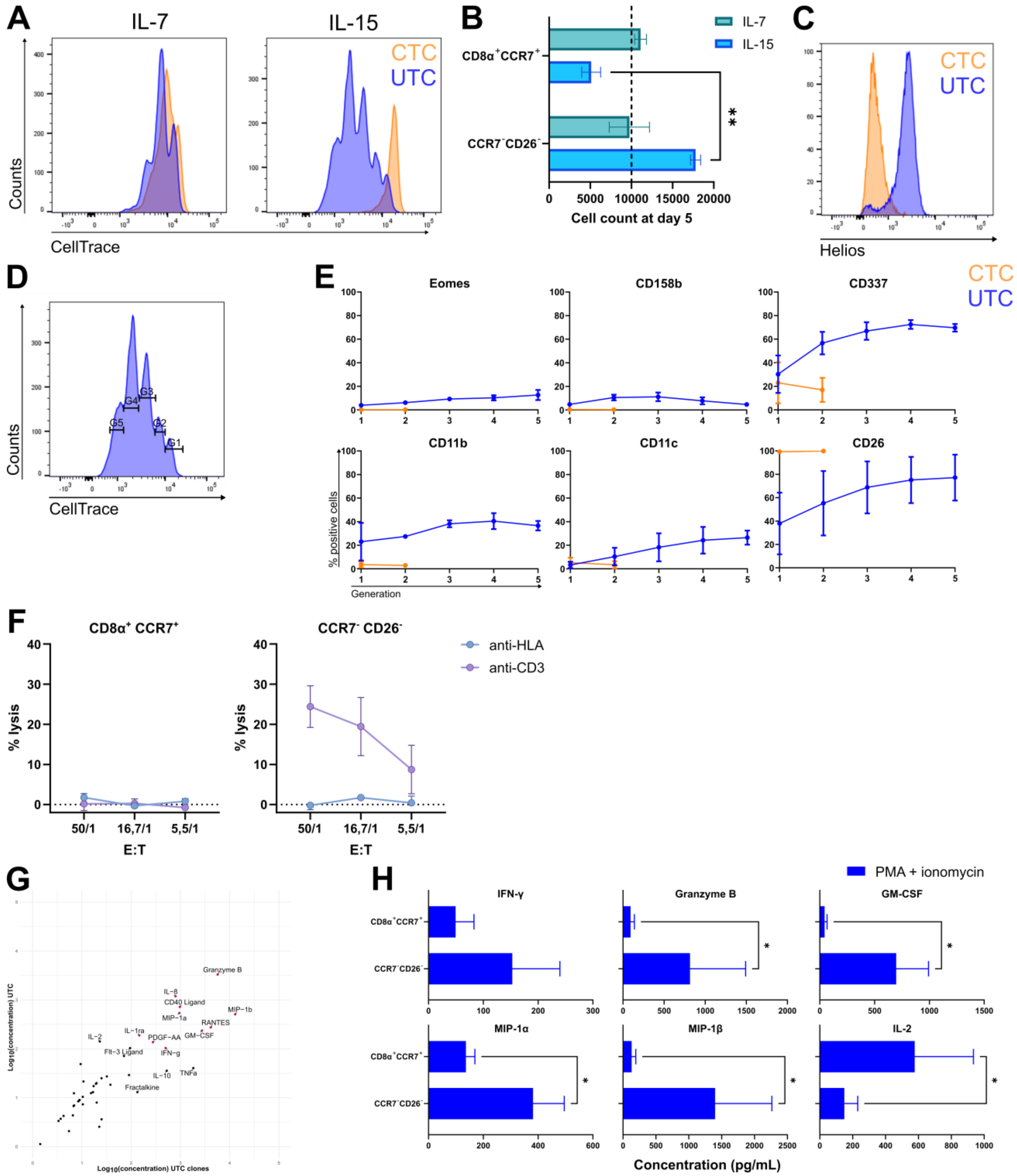

**Supplementary Fig. 5**. **UTC phenotype and functionality after proliferation with IL-15.** (**A**) Proliferation assessed by CellTrace Violet dye dilution for the CTCs (orange) and UTCs (blue) isolated from CB, after 5 days of incubation in the presence of IL-7 (10 ng/mL) or IL-15 (10 ng/mL). Representatives of three experiments. (**B**) Cell count in CTC and UTC cultures after five days of proliferation with IL-7 or IL-15, started at day 0 with 10 000 cells per population (dotted line) (mean ± SEM, n = 2). (**C**) Representative flow cytometric analysis of Helios expression by the (remaining) CTCs and UTCs, after five days of proliferation with IL-15. (**D**) Gating strategy for the five generations in figure S6E, indicating successive rounds of cell division. Proliferation visualized by

CellTrace Violet dye dilution from right to left. (**E**) Phenotypical changes of the CTCs or UTCs during proliferation with IL-7 or IL-15 respectively, measured using flow cytometry and plotted per generation (Figure S6D) (mean ± SEM, n = 2). (**F**) Cell lysis of anti-HLA (W6/32 hybridoma) or anti-CD3 (OKT3 hybridoma) target cells after 4 h of co-incubation CTCs or UTCs in different effector-target ratios (E:T). The CTCs and UTCs were sorted from CB and incubated overnight with IL-15, before the assay (mean ± SEM, n = 3). (**G**) Screening of cytokine secretion of 44 soluble cytokines (table S5) was measured using a multiplex immunoassay. $Log_{10}$-value of the secreted concentration by freshly sorted UTCs and UTC clones after 24 h of stimulation with PMA + ionomycin. Mean secretion of three UTC clones is shown. (**H**) Secreted IFN-γ, Granzyme B, GM-CSF, MIP-1α, MIP-1β and IL-2 measured in the supernatant of CTCs and UTCs after stimulation with PMA + ionomycin for 24 h. Wilcoxon matched-pairs signed rank test was used to assess statistically significant differences in cytokine secretion between the populations (mean ± SEM, n = 7). P-value < 0.05 (*).

**Supplementary table S1. Significantly differentially expressed between the PD-1$^+$ and PD-1$^-$ population, in both PNT and CB**

| *RNA* | *Protein* |
|---|---|
| TBKBP1 | RBP4 |
| SIPA1L2 | YWHAH |
| ZNF683 | GIMAP1 |
| RTKN2 | SPTBN1 |
| MME | |
| AC004585.1 | |
| TRGC2 | |
| PLCH2 | |
| TNFRSF9 | |
| FZD3 | |
| GNG4 | |
| FCRL3 | |
| TBX21 | |
| PTPN14 | |
| PDCD1 | |
| PON2 | |
| XCL1 | |
| RHOC | |
| VAV3 | |
| CXCR3 | |
| GCSAM | |
| DUSP2 | |
| ID3 | |
| RTP5 | |
| PIK3R1 | |
| LATS2 | |
| IKZF2 | |
| SYTL2 | |
| UBASH3B | |
| CHSY1 | |
| RTL6 | |
| RUNX3 | |
| SURF4 | |
| CNR2 | |
| TP53INP1 | |
| CHST2 | |
| TRGC1 | |
| ATP2B4 | |
| HELZ | |
| PRKX | |
| RTN4 | |
| TRG-AS1 | |
| NEAT1 | |
| P2RX5 | |

| |
|---|
| HEG1 |
| ITPR2 |
| TOX |
| CDC34 |
| ITGB2-AS1 |
| RAB37 |
| PFKP |
| SMC4 |
| PPP2R5C |
| FLNA |
| PRKCH |
| MAPKAPK2 |
| ERGIC1 |
| HERC1 |
| ELOVL5 |
| RGS2 |
| RAB8B |
| ELF1 |
| CLDND1 |
| CCDC69 |
| MKRN1 |
| S100A10 |
| LAPTM4A |
| TNRC6B |
| SORL1 |
| IRF2BP2 |
| SATB1 |
| STK17B |
| IL7R |
| PEBP1 |
| COTL1 |
| ATP1A1 |
| AAK1 |
| ARRB2 |
| BACH2 |
| RHOH |
| SOD1 |
| RFLNB |
| TMSB10 |
| PARK7 |
| IKZF1 |
| LDLRAP1 |
| FAM102A |
| TAGLN2 |
| CD2 |
| ITM2A |

| |
|---|
| AUTS2 |
| IRF2BPL |
| LBH |
| GNG2 |
| SNHG32 |
| LAPTM5 |
| CD3D |
| RASSF5 |
| SARAF |
| CCND2 |
| ABLIM1 |
| LINC00861 |
| RPS27 |
| PGGHG |
| CHI3L2 |
| RPLP2 |
| RPL27 |
| TPT1 |
| POU2F2 |
| RPL38 |
| RPS21 |
| RPL31 |
| AP1M1 |
| NDFIP1 |
| PAG1 |
| RPS8 |
| ULK2 |
| TPM2 |
| AIF1 |
| ARMH1 |
| GRAP2 |
| SELL |
| FOXP1 |
| RORA |
| KCNQ1 |
| GLRX |
| RCAN3 |
| CD8B |
| PASK |
| TRABD2A |
| CCR7 |
| CSGALNACT1 |
| LINC01550 |
| SSTR3 |

**Supplementary table S2. Cell number in each identified cluster**

| *Cluster* | *Number of cells* |
|---|---|
| MME+ UTC | 2080 |
| GNLY- MME- UTC | 3220 |
| IL32+ UTC | 989 |
| GNLY+ UTC | 2830 |
| GZMK+ DN UTC | 2563 |
| CTC 1 | 4104 |
| CTC 2 | 3053 |
| CTC 3 | 2300 |
| CTC 4 | 1316 |
| CTC 5 | 567 |
| Cycling T | 677 |
| NKT/MAIT | 816 |
| NK | 212 |

**Supplementary table S3. Top 10 differentially expressed genes for each identified cluster**

| *MME+ UTC* | *GNLY- MME- UTC* | *IL32+ UTC* | *GNLY+ UTC* | *GZMK+ DN UTC* | *CTC 1* | *CTC 2* | *CTC 3* | *CTC 4* | *CTC 5* | *Cycling T* | *NKT/MAIT* | *NK* |
|---|---|---|---|---|---|---|---|---|---|---|---|---|
| ID3 | DUSP2 | IL32 | GNLY | GZMK | S100B | CD8B | S100B | S100B | ACTG1 | LGALS1 | KLRB1 | GNLY |
| PPP2R5C | JUNB | GNLY | DUSP2 | LYAR | PASK | SOX4 | PASK | PASK | CCL5 | GZMA | ANXA1 | NKG7 |
| RTN4 | ID3 | NKG7 | NR4A2 | CCL5 | TOB1 | AIF1 | PRKCQ-AS1 | RPS8 | SELL | GZMK | RGCC | GZMB |
| ZNF683 | ZNF683 | CTSW | NKG7 | CMC1 | ANXA1 | ARMH1 | RPS8 | SNHG29 | PASK | LGALS3 | MT2A | CCL3 |
| SMC4 | ZFP36L2 | KLRC2 | ZFP36 | PLAC8 | NELL2 | CHI3L2 | NELL2 | RPS3A | CD27 | CD81 | TNFAIP3 | GZMH |
| MME | NR4A2 | DUSP2 | ZFP36L2 | ARID5B | RPL23 | SNHG29 | RPS3A | AIF1 | GAPDH | S100A4 | S100A4 | FGFBP2 |
| RTKN2 | ZFP36 | HOPX | XCL1 | GPR183 | USP10 | FHIT | RPS18 | RPS18 | SMC4 | ANXA2 | TOB1 | SPON2 |
| TRGC2 | DUSP1 | CST7 | PIK3R1 | JUN | ARMH1 | LINC01089 | RPS6 | RPS12 | FYB1 | STMN1 | LST1 | CCL4 |
| HELZ | KLF2 | ZNF683 | IER2 | CLDND1 | SELL | FOXP1 | SNHG29 | NOSIP | GZMK | HMGB2 | USP10 | KLRB1 |
| KDM5B | CTSW | PIK3R1 | ID1 | TXNIP | TNFAIP3 | RPL23 | ARMH1 | RPS4X | CARS1 | S100A6 | LTB | TYROBP |

**Supplementary table S4. CITE-seq cell surface protein antibody panel**

| | | | | | |
|---|---|---|---|---|---|
| CD80.1 | CD86.1 | CD274.1 | CD273 | CD275-A0009 | CD276.1 |
| CD11b-mh | Galectin9 | CD270 | CD252 | CD137L | CD155 |
| CD112 | AnnexinV | CD47.1 | CD70.1 | CD30 | CD48.1 |
| CD277 | CD40.1 | CD154 | CD52.1 | CD3-A0034 | CD4-A0045 |
| CD8 | CD56-A0047 | CD45-A0048 | CD3-A0049 | CD19.1 | CD14-A0051 |
| CD33.1 | CD11c | CD34.1 | CD138-A0055 | CD269 | B2M.1 |
| HLA-ABC | CD90-A0060 | CD117 | CD10 | CD45RA | CD123 |
| CD7.1 | CD105 | CD201 | CD49f | CD194 | CD4-A0072 |
| CD44-mh | CD15-mh | CD8a | CD14-A0081 | CD16 | CD56-A0084 |
| CD25 | CD45RO | CD279 | TIGIT.1 | IgG2a-Mouse-k | IgG2b-Mouse-k |
| IgG2b-Rat-k | CD20 | CD335 | CD294 | CD45R-B220 | CD326 |
| CD31 | CD44.1 | CD133-A0126 | Podoplanin | CD140a | CD140b |
| Cadherin | EGFR.1 | CD340 | CD146 | CD324 | IgM |
| CD5.1 | TCRgd | CD183 | CD195 | CD32 | CD196 |
| CD185 | CD103 | CD69.1 | CD62L | CD197 | CD161 |
| CD152 | CD223 | KLRG1.1 | CD27-A0154 | CD107a | CD95 |
| CD134 | HLA-DR | CD1c | CD11b | CD64 | CD141 |
| CD1d | CD314 | CD66b | CD35 | CD57 | CD366 |
| CD272 | CD278 | CD275-A0172 | CD58.1 | CD96.1 | CD39 |
| CD178 | CD24.1 | CD21 | CD11a | CD79b | CD66a-c-e |
| CD244.1 | CD27-A0191 | CD235ab | Siglec8 | CD206 | CD169 |
| CD370 | XCR1.1 | Notch-A0213 | Integrin | CD268 | CD42b |
| CD54 | CD62P | CD119 | IRF5.1 | ERK1 | RORg |
| TCRab | Notch-A0233 | CD68.1 | IgG1-Rat-k | IgG2a-Rat-k | IgG2a |
| IgG-Hamster | CD192 | CD102 | CD106 | CD122 | CD267 |
| CD62E | IRF4.1 | KLRG1-mh | CD135 | FceRIa | CD41 |
| CD137 | CD254 | CD43 | CD163.1 | CD83.1 | CD357 |
| CD59.1 | CD309 | CD13 | CD184 | CD2.1 | CD226-A0368 |
| CD29 | CD303 | CD49b | CD61 | CD81.1 | CD98 |

| | | | | | |
|---|---|---|---|---|---|
| IgG | CD177.1 | CD55.1 | IgD | CD18 | CD28.1 |
| TSLPR | CD38-A0389 | CD127 | CD45-A0391 | CD15 | CD22.1 |
| CD71 | B7H4 | CD26 | CD193 | CD115 | CD204 |
| CD144 | CD301 | CD1a | CD207.1 | CD63.1 | CD284 |
| CD304 | CD36.1 | CD172a | CD85g | CD38-A0410 | VSIG4.1 |
| CD243 | CD72.1 | CD158 | MERTK.1 | Folate | Tim4 |
| CD171 | CD230 | CD325 | ReceptorD4 | GABRB3.1 | CD207-mh |
| TMEM119.1 | CD93.1 | CD200.1 | CD338 | C5L2 | CD235a |
| CD49a | CD49d | CD73 | CD79a | CD9.1 | mastCellTryptase |
| TCRa7 | TCRd2 | TCRg9 | TCRa24 | CD354 | CD202b |
| CD305 | LOX1 | CD158b | CD203c | CD133-A0594 | CD209.1 |
| CD110 | CD158e1 | CD158f | CD337 | CD253 | CD186 |
| CD226-A0805 | CD205 | CD271 | CD109.1 | CD11a-CD18 | CD126 |
| CD164.1 | CD142 | CD307c-FcRL3 | CD307d | CD307e | CD319 |
| CD138-A0831 | CD199 | CD45RB | CD99.1 | CD371 | CD46.1 |
| CD151.1 | CD218a | CD257 | CLEC1B.1 | CD94 | IgE |
| CD365 | CD150 | CD162 | Ig-lightChainK | Mac2 | CD85j |
| CD23 | Ig-lightChainL | HLA-A2 | CD198 | GARP | CD4-A0922 |

**Supplementary table S5. Luminex Performance Human XL Cytokine Magnetic Panel 44-plex Fixed Panel**

| | |
|---|---|
| CCL2/JE/MCP-1 | IL-1ra/IL-1F3 |
| CCL3/MIP-1 alpha | IL-2 |
| CCL4/MIP-1 beta | IL-3 |
| CCL5/RANTES | IL-4 |
| CCL11/Eotaxin | IL-5 |
| CCL19/MIP-3 beta | IL-6 |
| CCL20/MIP-3 alpha | IL-7 |
| CD40 Ligand/TNFSF5 | IL-8/CXCL8 |
| CXCL1/GRO alpha/KC/CINC-1 | IL-10 |
| CXCL2/GRO beta/MIP-2/CINC-3 | IL-12 p70 |
| CXCL10/IP-10/CRG-2 | IL-13 |
| EGF | IL-15 |
| FGF basic/FGF2/bFGF | IL-17/IL-17A |
| Flt-3 Ligand/FLT3L | IL-17E/IL-25 |
| G-CSF | IL-33 |
| GM-CSF | PD-L1/B7-H1 |
| Granzyme B | PDGF-AA |
| IFN-alpha 2/IFNA2 | PDGF-AB/BB |
| IFN-beta | TGF-alpha |
| IFN-gamma | TNF-alpha |
| IL-1 alpha/IL-1F1 | TRAIL/TNFSF10 |
| IL-1 beta/IL-1F2 | VEGF |
